# Glycan-reactive antibodies isolated from human HIV-1 vaccine trial participants show broad pathogen cross-reactivity

**DOI:** 10.1101/2025.01.17.633475

**Authors:** Parker J. Jamieson, Xiaoying Shen, Alexandra A. Abu-Shmais, Perry T. Wasdin, Katarzyna Janowska, Robert J. Edwards, Garrett Scapellato, Simone I. Richardson, Nelia P. Manamela, Shuying Liu, Maggie Barr, Rebecca A. Gillespie, Jessica Mimms, Naveenchandra Suryadevara, Ty A. Sornberger, Seth Zost, Rob Parks, Shelby Flaherty, Alexis K. Janke, Bethany N. Howard, Yukthi P. Suresh, Ruth M. Ruprecht, James E. Crowe, Robert H. Carnahan, Justin R. Bailey, Kanekiyo Masaru, Barton F. Haynes, Penny L. Moore, Priyamvada Acharya, David C. Montefiori, Spyros A. Kalams, Shan Lu, Ivelin S. Georgiev

## Abstract

HIV-1 continues to pose a significant global health challenge, requiring ongoing research into effective prevention and treatment strategies. Understanding the B cell repertoire that can be engaged upon vaccination in humans is crucial for the development of future preventive vaccines. In this study, PBMCs from HIV-negative participants in the multivalent HVTN124 human HIV-1 vaccine clinical trial were interrogated for HIV-reactive B cells using LIBRA-seq, a high-throughput B cell mapping technology. We report the discovery of glycan-reactive antibodies capable of neutralizing diverse heterologous HIV-1 virus strains. Further, isolated antibodies showed broad cross-reactivity against antigens from a variety of other pathogens, while remaining mostly negative on autoreactivity assays. The emerging class of glycan-reactive virus-neutralizing antibodies with exceptional breadth of pathogen cross-reactivity may present an effective target for vaccination at the population level.

## Importance

Understanding how the human immune system recognizes and combats viruses is crucial for developing better vaccines and treatments. Here, through characterization of the B cell receptor repertoires of participants in HVTN124, a multivalent HIV-1 vaccine human clinical trial, we discovered antibodies that recognize sugar molecules (glycans) on antigens from a range of unrelated viral families, yet do not show reactivity toward self-antigens. In addition to their binding breadth, these antibodies can also neutralize multiple diverse strains of HIV-1. Our findings reveal an emerging and underappreciated mechanism for antibodies to counteract virus infection, potentially opening doors for developing vaccines that preferentially elicit glycan-reactive antibody species to broadly protect against different viruses.

Human immunodeficiency virus type 1 (HIV-1) is a positive-sense ssRNA retrovirus that affects 38.4 million people globally and is the causative agent for acquired immunodeficiency syndrome (AIDS). Once transmitted via bodily fluids, HIV-1 will target CD4+ cells and integrate into the host cell genome. Integration allows for the constant production of new HIV-1 virions; however, this is met by a strong adaptive immune response to reduce the total viral load. In this tug-of-war between HIV-1 and the immune system, HIV-1 can create viral escape mutants that increase viral diversity and the potential to evade the host’s adaptive immune response [1–7]. A common source of escape exists in the surface viral envelope (Env) gene, which encodes the gp160 envelope glycoprotein trimer. The gp160 trimer allows HIV-1 to target, bind, and enter CD4+ cells. The gp160 trimer is divided into two subunits: a gp120 receptor/co-receptor-binding protein and a gp41 transmembrane protein [1–7]. The gp120 subunit comprises critical epitopes that influence HIV-1 infectivity and are primary antibody targets. Some of these epitopes include the CD4 binding site for initial binding to CD4+ cells, the V1/V2 loop for trimer stabilization, and the V3-glycan loop for co-receptor recognition. Because the gp160 trimer is exposed on the viral surface, it faces constant selective pressure by the adaptive immune system, resulting in frequent mutations that confer antibody evasion. In addition to sequence diversity, HIV-1 Env is comprised of a host-derived glycan shield that can mask critical epitopes from neutralizing antibodies. By mutating Env and other viral genetic components, along with displaying self-like carbohydrate structures, HIV-1 can diversify into distinct groups, clades, and present different gp160 epitopes, thus creating a major obstacle in regard to HIV-1 vaccine development [1–7].

While effective vaccines, such as the mRNA COVID-19 vaccine, have mitigated many infectious diseases, there is still no effective preventive HIV-1 vaccine. Antiretroviral medications suppress viral replication effectively; however, individuals infected with HIV- 1 continue to harbor the virus and have to take antiretroviral therapies (ART) throughout their lives, and are subjected to long-term drug toxicity. The most promising HIV-1 vaccine trial to date was the RV144 trial in Thailand, which tested a canarypox vector prime-Env gp120 protein boost vaccine. However, the 31.2% efficacy of RV144 was not robust, and subsequent canarypox vector-Env gp120 protein vaccine trials in South Africa failed to show efficacy [8–16]. The development of a safe and protective vaccine against HIV-1, though a formidable challenge, remains a worldwide health priority.

A challenge for HIV-1 vaccine design is eliciting effective HIV-1 antibodies that protect against a diverse range of viral variants. HVTN124, developed by Worcester HIV Vaccine, was a phase 1A human vaccine trial that was completed in October 2020 and utilized a second-generation polyvalent DNA–protein HIV vaccine consisting of DNA inserts expressing four HIV-1 gp120 antigens and an HIV-1 gag gene, and the four matching adjuvanted recombinant HIV-1 gp120 proteins in both prime–boost and coadministration regimens aiming to elicit high magnitude, cross-reactive antibodies against multiple HIV-1 clades [17]. Immunogens used included HIV-1 Env gp120 DNA and proteins from clades A (92UG037.8), B (JR-FL), C (93MW965.26), and AE (ConAE (Ref: 92KDSu5-19150)), along with DNA of the conserved HIV-1 Gag polyprotein and a glucopyranosyl lipid adjuvant in a stable emulsion (GLA-SE) to stimulate a Th1-dominant response [18–21]. The HVTN124 clinical trial included a total of 60 HIV-negative men and women across different age ranges and ethnic backgrounds. Participants were randomly divided into three groups: Group 0 for establishing safety, Group 1 for staggered immunizations, and Group 2 for concurrent immunizations. A schematic of the HVTN124 immunization strategies, groups, and composition is depicted in Figure 1A. Preclinical experiments in rhesus macaques, mice, and rabbits identified that staggered immunizations (Group 1 approach) elicited antibodies with the highest avidity, high titers, and broad reactivity [22–27]. In one of these preclinical animal studies, the HVTN124 vaccine composition and staggered immunizations in rabbits elicited high IgG titers against the V1/V2 epitope, which is responsible for trimer stabilization, neutralization resistance, and modulates viral entry [27]. In the HVTN124 trial, Groups 1 and 2 both elicited polyclonal cross-reactive HIV-1 antibodies in serum; however, the monoclonal antibody response had yet to be characterized for either group. Monoclonal antibody characterization is essential for HIV-1 vaccine design as it informs features of antibodies that may help define their effectiveness. These include the ability of a vaccine to elicit antibodies that recognize diverse epitopes across inter- and intra-clade HIV-1 strains, or cross-react, and neutralize the virus.

**Figure 1.**
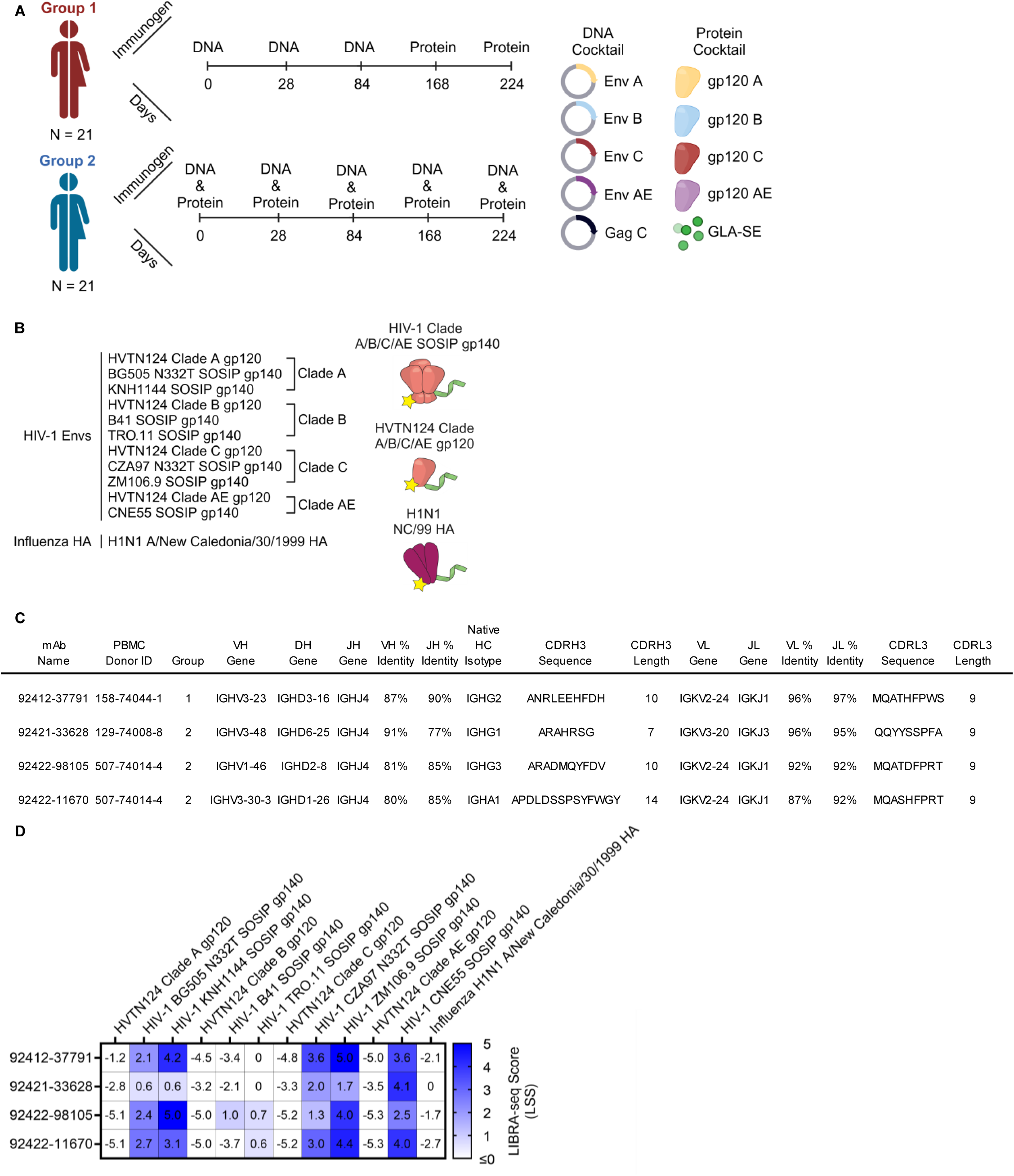
LIBRA-seq discovery of mAbs from HVTN124 donor PBMCs. (A) HVTN124 vaccine trial. Graphic of the HVTN124 immunization groups, timepoints, and makeup. Figure was created using BioRender. (B) Antigen screening library used to interrogate the B cell repertoire from HVTN124 donor PBMCs. The library consisted of four HVTN124 gp120 immunogens, seven SOSIP gp140 antigens, and one influenza HA antigen. Figure was created using BioRender. (C) Sequence features of identified HIV-1 Env-reactive mAbs isolated from HVTN124 donor PBMCs. PBMC donor identity and vaccine group are also provided. All antibodies were expressed as IgG1 for further characterization. (D) Antigen-specific LIBRA-seq scores for the four HIV-1 Env-reactive mAbs (rows) are shown for the antigens used in the screening library (columns). Heatmap of LIBRA-seq scores ranges from white (low) to blue (high).

Here, we report the isolation and characterization of a set of glycan-reactive, including Fab-dimerized glycan-reactive (FDG), antibodies from HVTN124 trial participants that broadly neutralize HIV-1 and hepatitis C virus (HCV). FDGs, such as the potent 2G12 broadly neutralizing antibody (bNAb) that utilizes a unique V_H_ domain swapping configuration resulting in Fab dimerization that presents as I-shaped, along with others such as DH851 that Fab dimerize without V_H_ domain swapping, have been previously reported in the literature as targeting conserved glycan patches [28]. A subset of these antibodies was found to be Fab-dimerized by negative-stain electron microscopy (NSEM) analysis. Moreover, some of these antibodies share public sequence signatures, or recurring complementarity-determining region (CDR) sequence motifs, with other known glycan-reactive antibodies, suggesting that such a public class of antiviral antibody recognition can be used as a target for broadly reactive vaccine candidates against HIV-1. We demonstrate that of the four antibodies isolated from HVTN124 PBMCs that bind to N-linked glycans, three of the four neutralized HIV-1, with one showing weak but broad neutralization across HIV-1 viruses from tiers 1 and 2. Further, we show that these antibodies cross-react with antigens from multiple diverse pathogens, including neutralization against HCV, but are mostly negative on autoreactivity assays. Overall, we propose that the elicitation of glycan-reactive antibodies can serve as an effective strategy for turning the elusive HIV-1 glycan shield into a Trojan horse, and therefore warrants further pursuit and optimization in future HIV-1 vaccine trials.

## Results

### Isolation of HIV-reactive antibodies from HVTN124 participants

To identify HIV-reactive B cells from HVTN124 participants, we performed LIBRA-seq, a high-throughput single-cell sequencing technology [29]. The LIBRA-seq antigen screening panel consisted of the four autologous HVTN124 gp120 vaccine immunogens (HVTN124 gp120 A, HVTN124 gp120 B, HVTN124 gp120 C, and HVTN124 gp120 AE) and seven heterologous SOSIP gp140 antigens (BG505, KNH1144, B41, TRO.11, CZA97, ZM106.9, and CNE55). Influenza H1N1 A/New Caledonia/30/1999 hemagglutinin (HA) was also included in the antigen screening panel as a negative control. HIV-1 SOSIP gp140 antigens were stabilized by an intermolecular disulfide bond between gp120 and gp41, displayed a truncated gp41 transmembrane region, and contained a variable flexible linker between gp120 and gp41 [30–35]. This 12-antigen panel, shown in Figure 1B, was applied separately to individual PBMC samples from three donors, resulting in the isolation of B cells with varying degrees of predicted cross-reactivity between multiple HIV-1 antigens. Notably, several B cell receptors (BCRs), isolated from multiple different HVTN124 participants, showed cross-reactivity between multiple HIV-1 antigens, with stronger signals for the SOSIP variants compared to the gp120 vaccine antigens, and binding was also observed for the influenza HA antigen (Figure 1C, Figure 2, and Figure S1). The preferential binding to SOSIP vs. gp120 vaccine antigens was in accordance with LIBRA-seq scores for the respective antigens (Figure 1D). Given the intriguing cross-reactivity with the influenza H1N1 A/New Caledonia/30/199 hemagglutinin (HA) antigen, we characterized these antibodies further.

**Figure 2.**
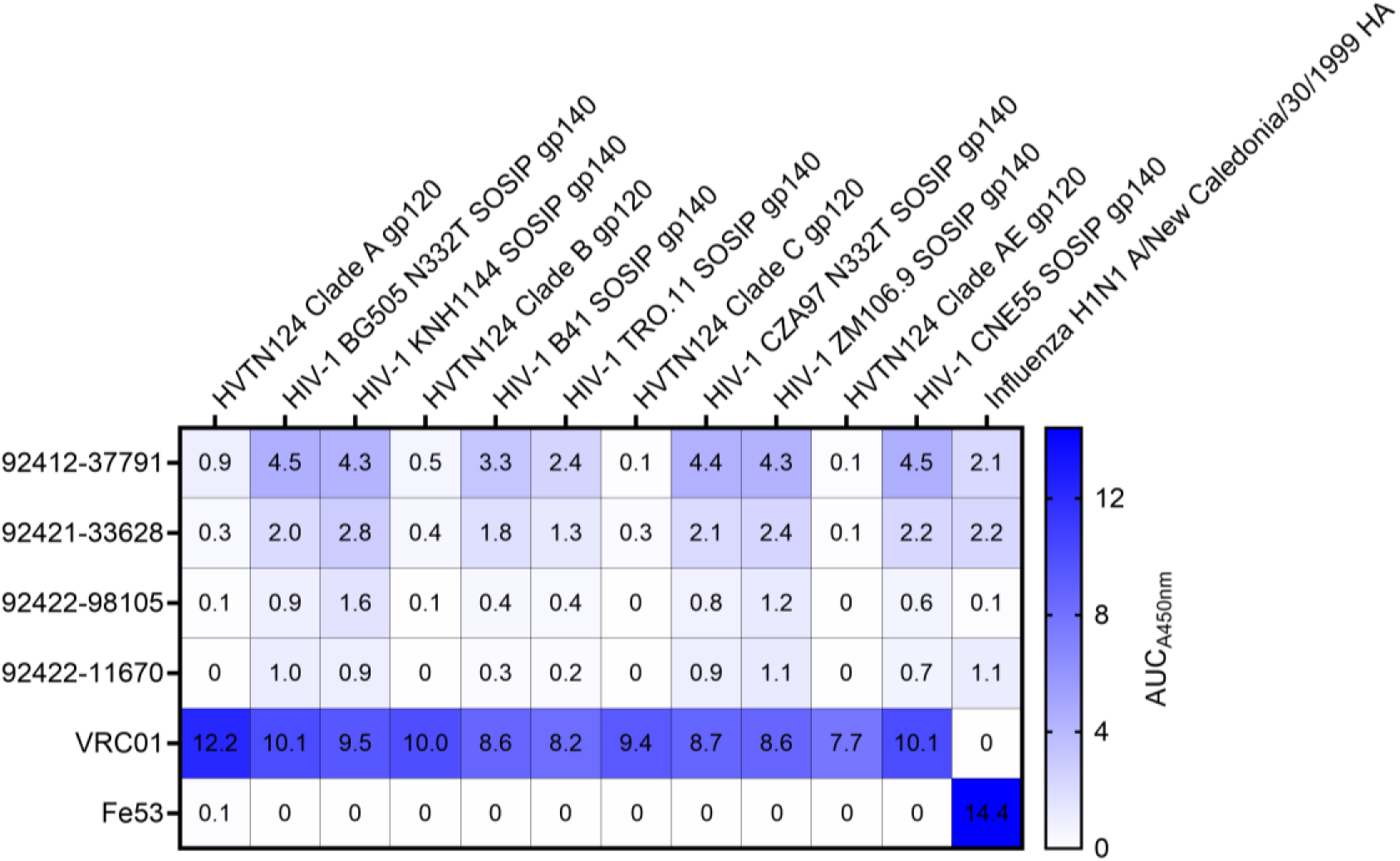
Characterization of HVTN124 mAbs by ELISA. ELISA validation of HVTN124 mAbs displayed as area under the curve (AUC) for the 12-antigen screening panel (columns) versus antibodies (rows), including the four HVTN124 mAbs, VRC01 (HIV-1 positive control), and Fe53 (influenza positive control). Values for all conditions were gathered in duplicate from a set of three repeats.

### HVTN124 antibodies target N-linked glycans on the surface of HIV-1 Env

After observing strong reactivity to influenza H1N1 A/New Caledonia/30/1999 HA by ELISA, we sought to identify what common features exist between HIV-1 Env and H1N1 Influenza HA. Since both antigens are glycoproteins, we elected to determine if these antibodies may target N-linked glycans as a common epitope. N-linked glycosylation sites on HIV-1 Env comprise roughly 50% of its molecular mass, with 70-90% of glycans on Env being forms of high mannose [36–38]. These host-derived N-linked glycans on Env have been established to form a glycan shield that helps HIV-1 evade NAbs by masking potentially neutralizing epitopes, along with ensuring proper Env folding and infectivity [39]. One of the most mannose-dense domains of HIV-1 Env lies at the third variable loop, or V3-glycan [40]. The V3-glycan is crucial for HIV-1 Env tropism and binding to the CCR5 or CXCR4 coreceptors, making conserved domains of the V3- glycan one of the primary targets for human glycan-reactive HIV-1 neutralizing antibodies [39, 41–44]. Seasonal influenza HA glycoproteins, which include H1N1, H3N2, and Victoria lineages, have been found to contain roughly 28-50% of high-mannose glycans; however, the proportion of high-mannose glycans on influenza HA can vary between strains [45]. Since HIV-1 Env and influenza HA both contain high-mannose glycans, we sought to test for antibody competition against D-(+)-Mannose, serving as a common glycan target between both viral antigens. As seen in Figure 3A and Figure S2, when incubated with 1M D-(+)-Mannose, all four of the HVTN124 mAbs and 2G12 showed a noticeable decrease in binding to HIV-1 CNE55 SOSIP gp140 by ELISA, whereas the V1/V2-reactive PG9 antibody, which recognizes gp120-associated high-mannose glycans but will not bind to protein-free glycans, showed no significant decrease in reactivity toward HIV-1 CNE55 SOSIP gp140 [46].

**Figure 3.**
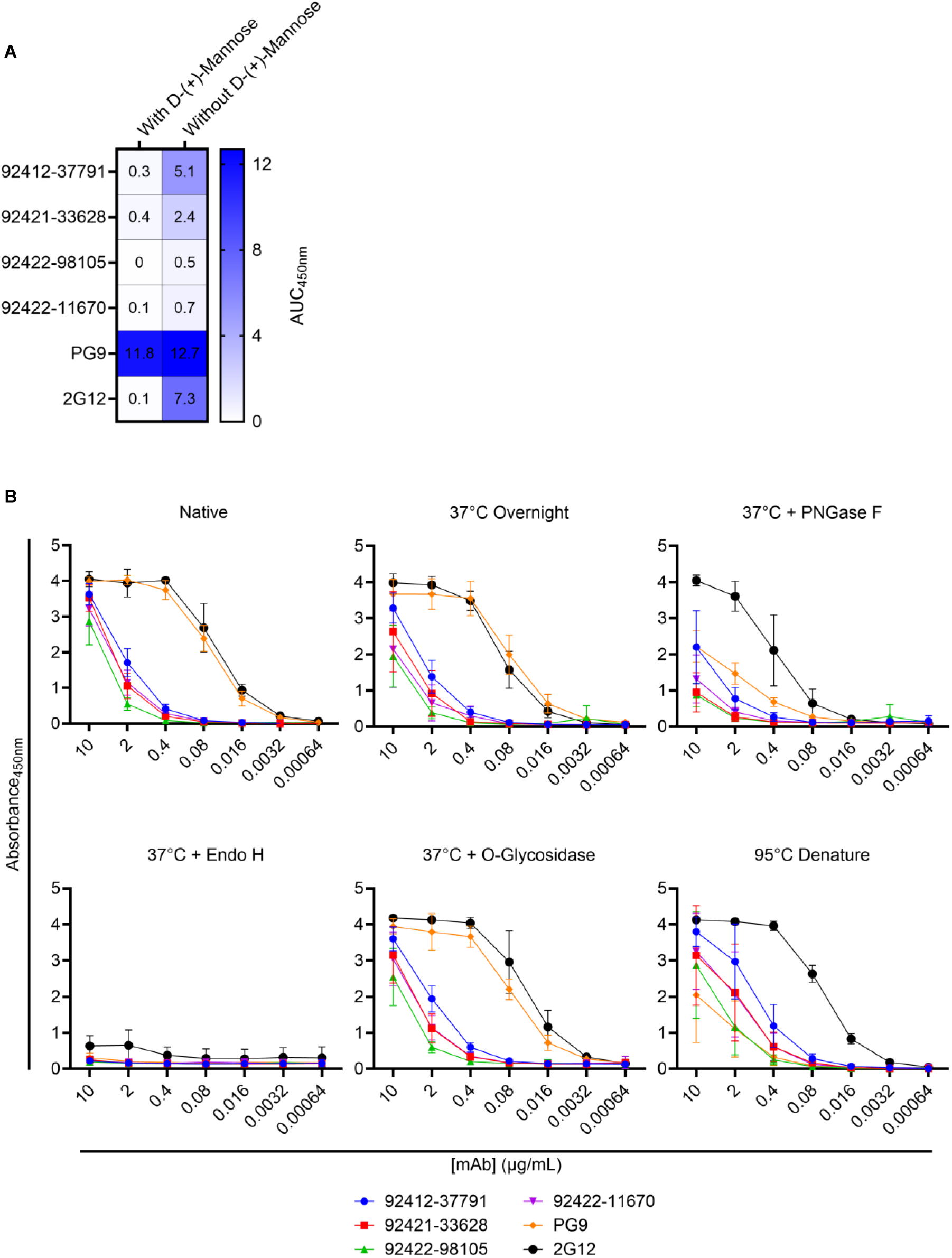
HVTN124 mAbs achieve broad reactivity via N-linked glycan recognition. (A) Antibody binding with and without 1M D-(+)-Mannose competition is displayed as a heatmap. The four HVTN124 mAbs, along with the V3-glycan-reactive 2G12 and V1/V2- reactive PG9 control mAbs (rows), were incubated with and without 1M D-(+)-Mannose against CNE55 (columns). Results are displayed from a set of three repeats in duplicate. (B) Enzymatic deglycosylation of HIV-1 CNE55 SOSIP gp140 tested by ELISA. The glycan-dependency of the HVTN124 mAbs was further evaluated by treating HIV-1 CNE55 SOSIP gp140 with different deglycosylation and/or denaturing conditions, then testing by ELISA. X-axis lists the antibody concentration in µg/mL, while absorbance at 450nm is listed along the Y-axis. Values for all six conditions are displayed from a set of three repeats in duplicate.

To further validate that these four HVTN124 mAbs bind to N-linked glycans, we performed a series of enzymatic deglycosylation reactions against HIV-1 CNE55 SOSIP gp140. Specific conditions of HIV-1 CNE55 SOSIP gp140 included: native, 37°C overnight, 37°C overnight with PNGase F, 37°C overnight with Endo H, 37°C overnight with O-glycosidase, and denatured at 95°C for 10 minutes. While the enzymes PNGase F and Endo H both cleave N-linked glycans from glycoproteins, PNGase F removes all high-mannose, complex, and hybrid N-linked glycans, whereas Endo H only removes high-mannose and some hybrid N-linked glycans. It has been previously reported that when CHO-cell-expressed HIV-1 gp120 protein is treated with Endo H, the binding affinity of 2G12 is significantly reduced due to the removal of high-mannose carbohydrates [47]. O-glycosidase only removes desialyated core 1 and core 3 O-linked disaccharides that are attached to Serine or Threonine residues. While not as extensive as N-linked glycans, O-linked glycans have been previously reported on a subset of HIV-1 isolates with unusually long V1 domains and can aid in shielding HIV-1 from V3- glycan bNAbs [48].

As shown in Figure 3B, the PNGase F and Endo H N-linked deglycosylation conditions displayed a noticeable reduction in antibody binding activity to HIV-1 CNE55 SOSIP gp140. Since HIV-1 can display O-linked glycans on the Env surface glycoprotein, we also opted to include the O-glycosidase O-linked deglycosylation condition; however, as shown in Figure 3B, there was no noticeable change in binding levels toward the HIV-1 CNE55 SOSIP gp140 with O-linked glycans removed. The 95°C denaturing condition was included to address if proper protein structure is necessary for antibody binding, which was shown to not be the case for each of the HVTN124 mAbs. These data indicate that the four HVTN124 mAbs are reactive to N-linked glycans on HIV-1 Env, specifically D-(+)-Mannose.

### Some HVTN124 antibodies do not show signs of autoreactivity

Next, we evaluated whether the four glycan-reactive antibodies display autoreactivity toward a panel of self-antigens. Three of the four antibodies showed no signal for any of the antigens in a standard AtheNA autoantigen panel of nuclear proteins, with only antibody 92412-37791 displaying any degree of autoreactivity for Centromere B and Histone (Figure 4A and Figure S3). When tested against cardiolipin, none of the four glycan-reactive antibodies displayed reactivity (Figure 4B and Figure S4). Next, we screened for reactivity against HEp-2 cells, which are used to identify autoantibodies, and observed that three of the four antibodies showed negligible to no signal; however, a positive signal was observed for one antibody, 92422-11670. Together, these results indicate that two of the four antibodies show no signs of autoreactivity, while the other two antibodies show diverse, yet minor, autoreactivity profiles.

**Figure 4.**
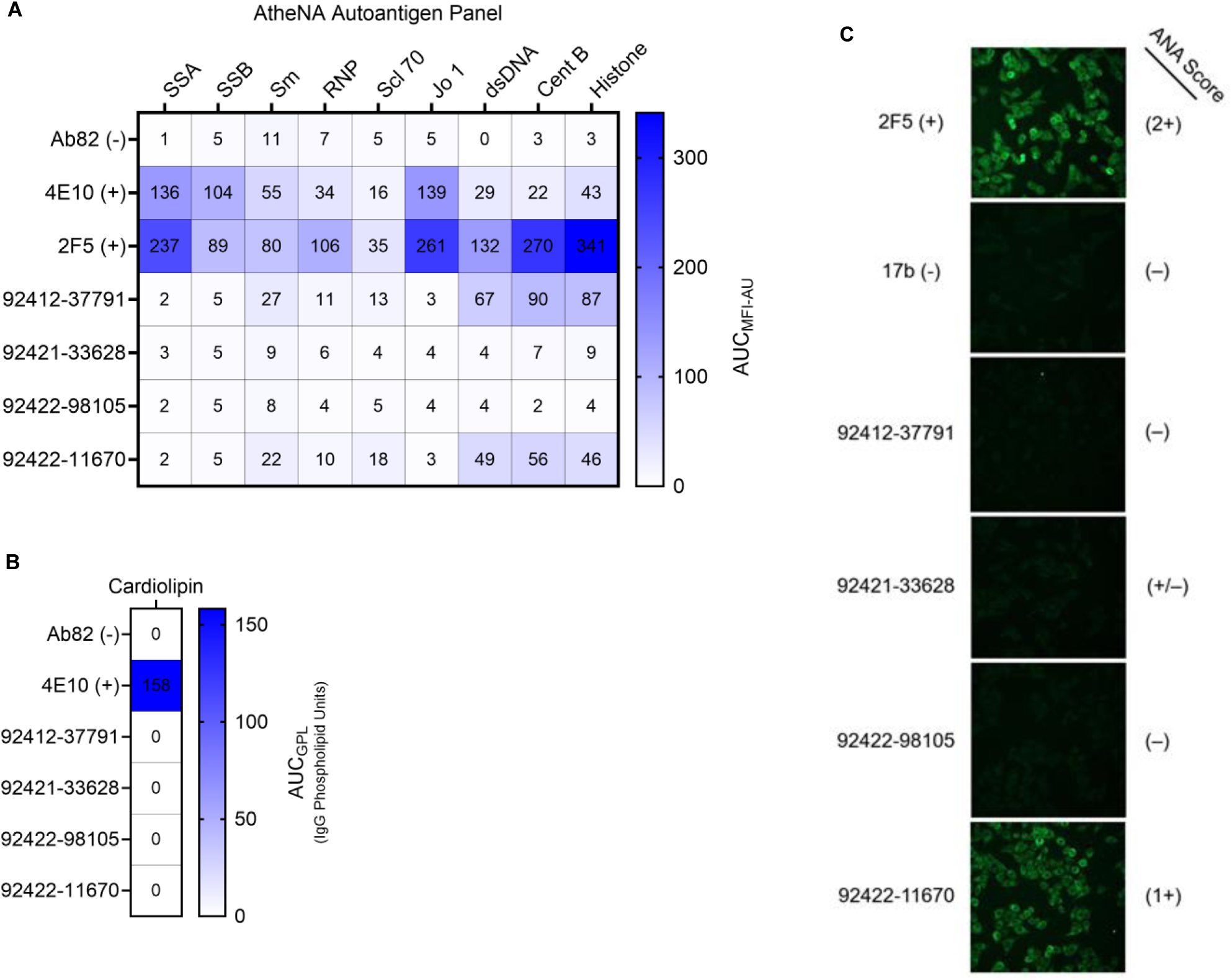
Autoreactivity analysis of HVTN124 mAbs. (A) Autoantigen reactivity against the AtheNA panel. Autoantigens are listed as columns, while antibodies are listed as rows. Positive control antibodies included 4E10 and 2F5, while Ab82 was used as a negative control antibody. In reference to Figure S3, values exceeding 120 MFI at 25 µg/mL for the AtheNA assay are considered positive. Values for all conditions are displayed from a set of two repeats. (B) Autoantigen reactivity against cardiolipin. Antibodies are listed as rows. 4E10 was used as a positive control antibody, while Ab82 was used as a negative control antibody. In reference to Figure S4, values at or greater than 20 GPL (IgG Phospholipid Units) at 50 µg/mL are considered positive. Values for all conditions are displayed from a set of two repeats. (C) HEp-2 cell monolayer culture immunofluorescence. HVTN124 mAbs were applied to whole, unpermeabilized, uninfected HEp-2 cells at 25 μg/mL mAb. Antibodies were detected using goat anti-human IgG-FITC (green). Positive and negative control mAbs used include 2F5 and 17B, respectively. 2F5, 17B, and 92422-11670 were all imaged at an exposure time of 5 seconds, while 92412-37791, 92421-33628, and 92422-98105 were imaged at an exposure time of 3 seconds. ANA scores are provided on the right. Images are at 40x magnification and were gathered in triplicate.

### A subset of HVTN124 antibodies display Fab-dimerization

It has been previously reported that glycan-reactive antibody species can be presented in a Fab-dimerized I-shaped conformation [28]. In HIV-naive humans, FDGs have been shown to be encoded by diverse antibody variable genes, with IGHV3-23 (24.3%), IGHV1-46 (18.9%), IGKV1-5 (10.8%), and IGKV2-24 (45.9%) being among the most frequent [28]. Interestingly, three of the four HVTN124 mAbs also utilized the IGKV2-24 light chain germline gene. To assess whether the HVTN124 mAbs are FDGs, we imaged the four HVTN124 mAbs as full IgG1 by negative stain electron microscopy (NSEM). We observed a heterogeneous mix of I- and Y-shaped full IgGs for 92412- 37791 and 92422-98105, while no I-shaped forms were observed for 92421-33628 and 92422-11670 (Figure 5). The Fab dimerized configuration of the two I-shaped mAbs were most reminiscent of the previously published DH1005 antibody [28].

**Figure 5.**
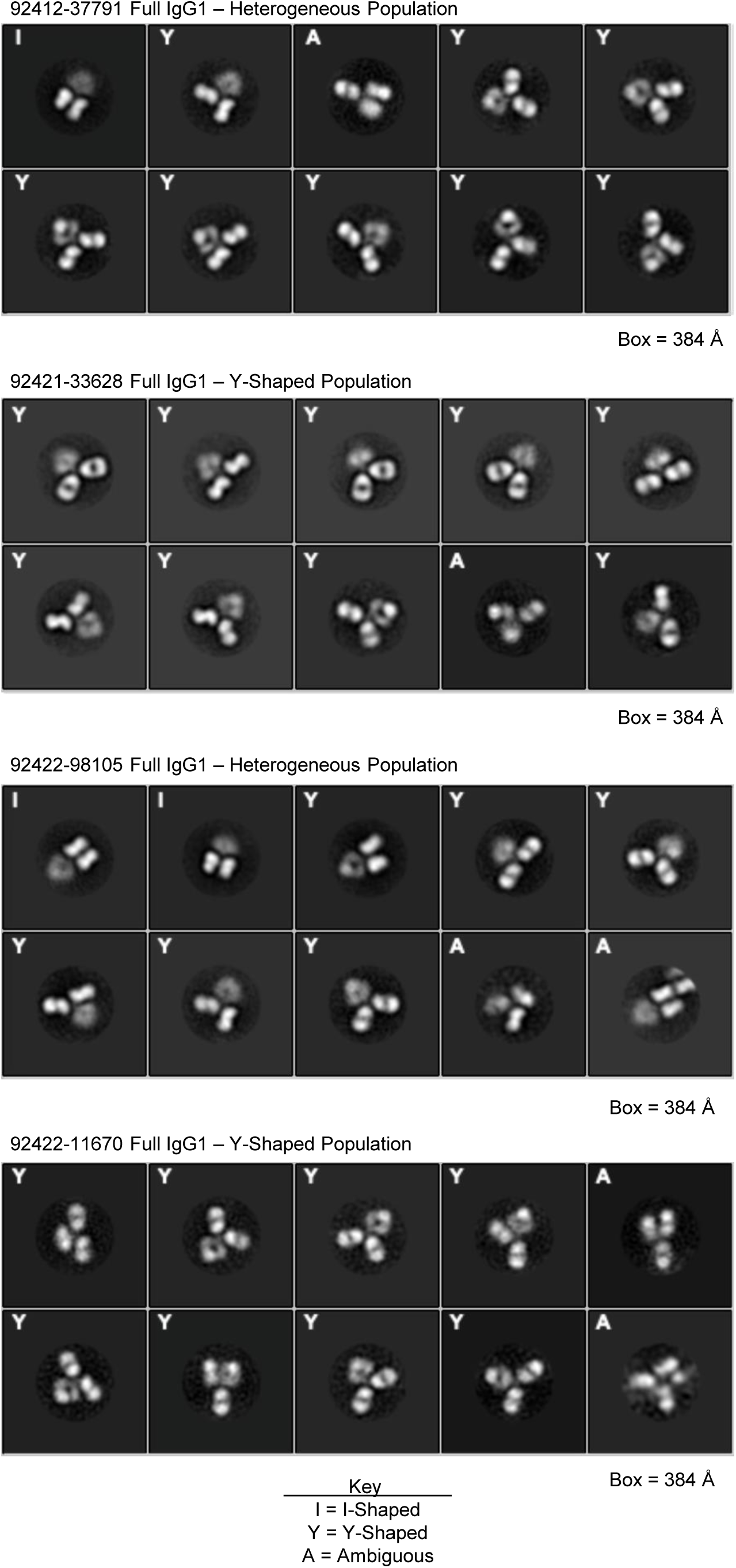
Negative stain electron microscopy. Negative stain electron microscopy 2D class averages of full IgG1 of 92412-37791 (top), 92421-33628 (2^nd^ from top), 92422-98105 (2^nd^ from bottom), and 92422-11670 (bottom). Imaging revealed that 92412-37791 and 92422-98105 are a heterogeneous population of I- and Y-shapes, while 92421-33628 and 92422-11670 are only Y-shaped. Images are ordered from most (top left corner) to least (bottom right corner) frequently observed by NSEM.

### Broad reactivity against multiple diverse viral antigens

Following the observation that the HVTN124 glycan-reactive antibodies broadly react to HIV-1 Env and influenza H1N1 A/New Caledonia/30/1999 HA, we evaluated whether these antibodies could recognize a broader antigen panel from a variety of other viruses. The 22 antigens used in this breadth panel were from HIV-1 (4 SOSIP gp140), influenza (2 group 1 HA, 5 group 2 HA, and 1 type B HA), coronavirus (6 SARS-CoV-2 spike variants and MERS-CoV spike), human parainfluenza virus type 3 (HPIV3) F0, hepatitis C virus (HCV) E1E2, and human cytomegalovirus (HCMV) gB (Figure 6). Each of the four HVTN124 mAbs showed broad reactivity against each of the viral families, albeit with variable binding strengths (Figure 6). The antigens in this breadth panel contain a different numbers of N-linked glycan sites (on average: 26-30 (HIV-1 Env), 5- 11 (influenza HA), 22 (SARS-CoV-2 spike), 5-6 (PIV3 F), 4-5 (HCV E1E2), and 18-19 (CMV gB), suggesting that these antibodies can target viral antigens with diverse glycan shield densities [49–54].

**Figure 6.**
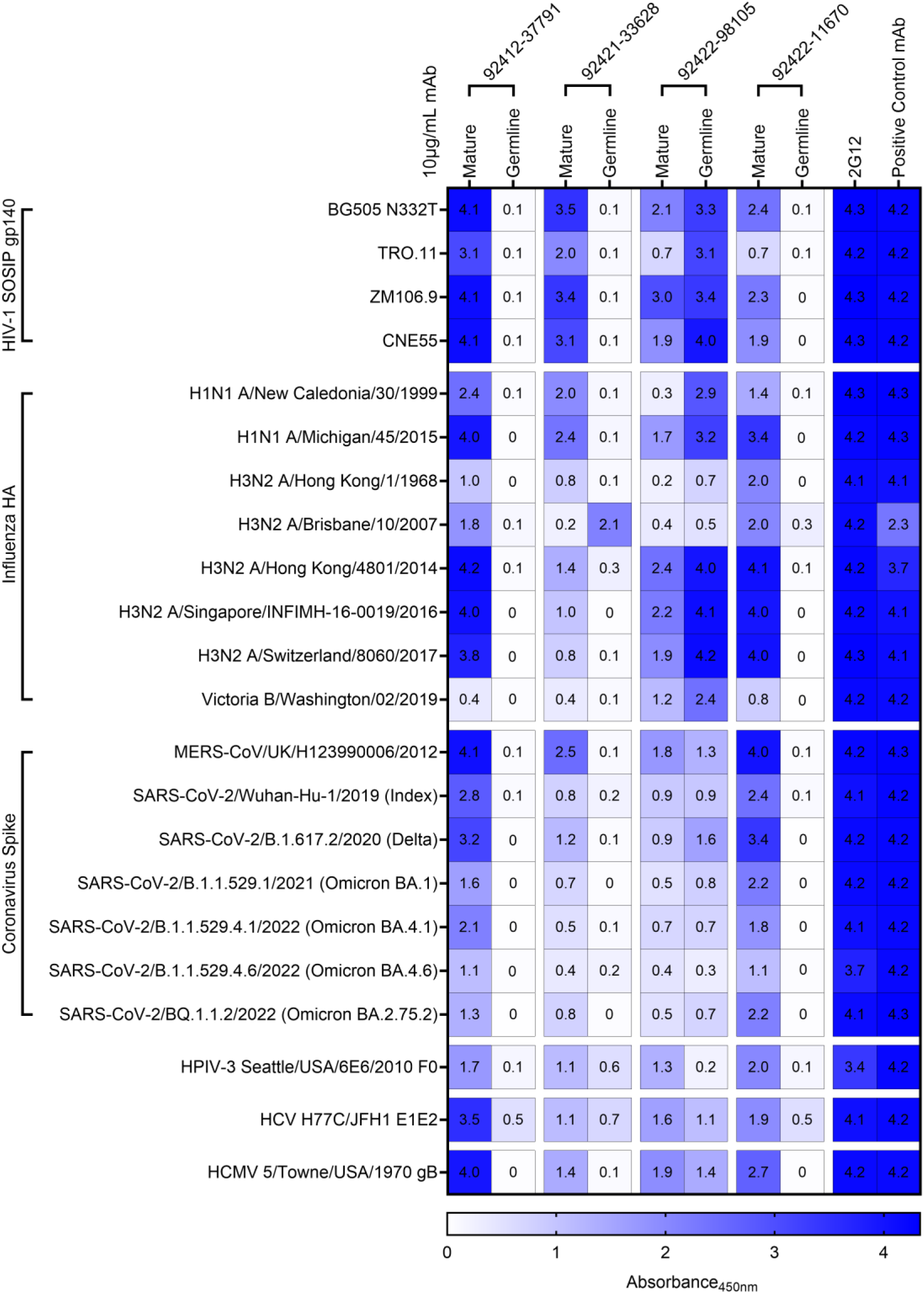
Expanded breadth analysis. HVTN124 mature and germline IgG1 mAbs vs. 22-antigen breadth panel. Four HVTN124 mAbs, 2G12, and the control mAbs are listed as columns, with the antigens used being listed as rows. Heatmap of absorbance values at 450nm (A450nm) using 10 µg/mL mAb is listed on the right from 0 in white to 4 in dark blue, with larger values in blue correlating to stronger antibody-antigen interactions. Control mAbs for each viral family include: VRC01 = HIV-1, CR9114 = influenza, 54043-5 = MERS-CoV and SARS- CoV-2, 3X1 = HPIV3, AP33 = HCV, IG2 = HCMV. ELISA 450nm absorbance values for all conditions in duplicate are displayed.

Next, we explored whether this level of broad reactivity was retained in germline-reverted versions of these antibodies, where both the heavy and the light chains for each antibody were reverted to their respective germline sequences apart from the CDR3 regions. Notably, the germline-reverted versions for three of the four antibodies showed virtually no or, in a small set, significantly reduced reactivity against the antigens tested (Figure 6). The only exception was the germline-reverted version of antibody 92422-98105 that exhibited strong recognition toward several of the antigens in the panel, similar to or, in some cases, even better than the mature antibody (Figure 6). Together, these results indicate that the breadth of antigen recognition for three of the four glycan-reactive antibodies is associated with acquired somatic hypermutation.

### Publicness of HVTN124 glycan-reactive antibody sequences

Sequences of the four HVTN124 mAbs were compared against a panel of known HIV-reactive antibodies, as well as against a panel of publicly available antibody sequences (Figure 7). Notably, three of the four HVTN124 mAbs exhibited substantial sequence similarity to the previously described glycan-reactive antibody DH1005 (Figure 7A). Antibodies 92412-37791 and 92422-11670 are encoded by the same light chain germline gene as DH1005, with 67% and 78% identity in the CDRL3 sequence, respectively. Further, antibody 92422-98105 is encoded by both the same heavy and light chain germline gene as DH1005, with CDRH3 amino acid identity of 60% and CDRL3 identity of 78%. Together, these results suggest that these HVTN124 mAbs may belong to a public light chain-driven class of glycan-reactive antibodies that can recognize HIV-1.

**Figure 7.**
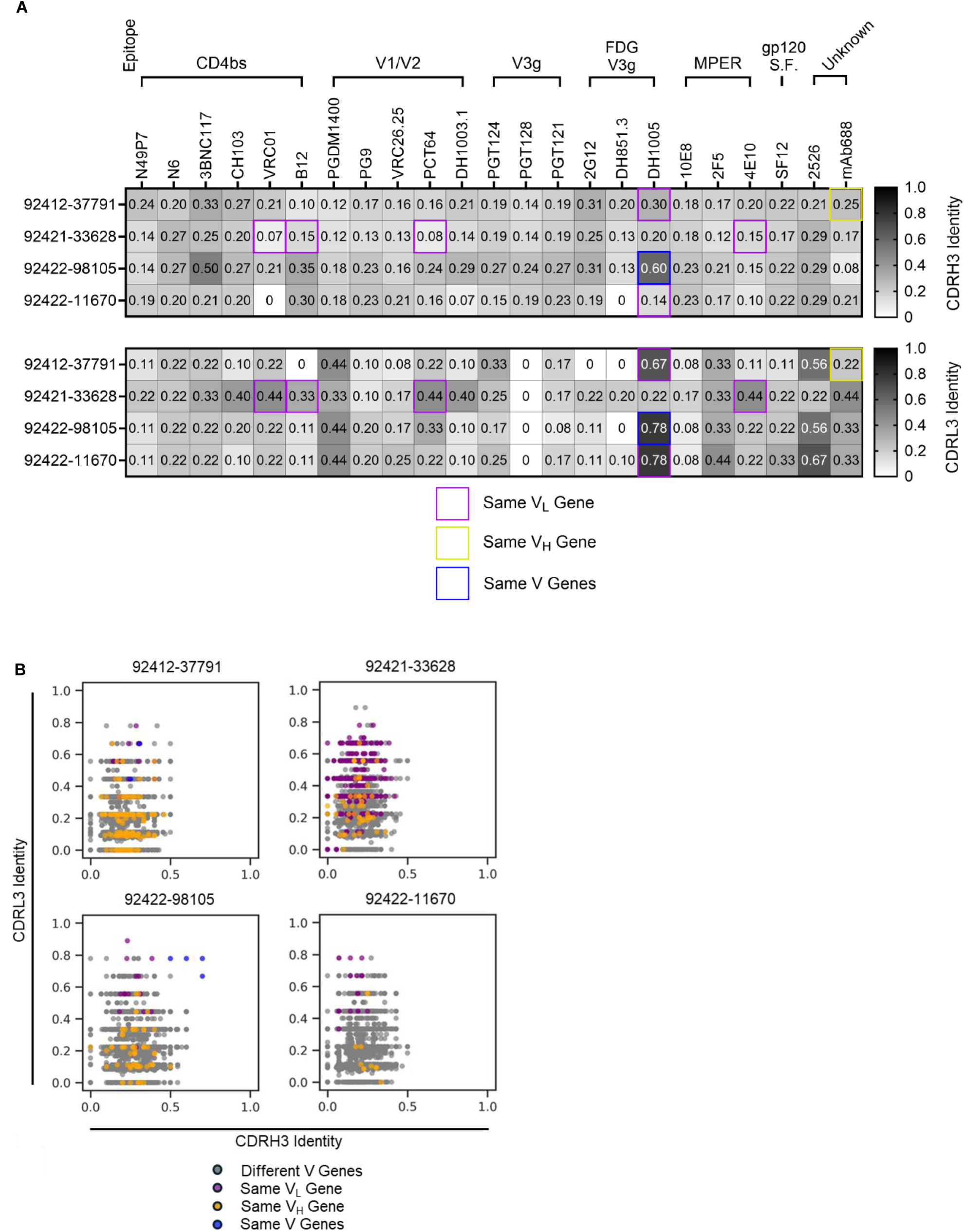
CDRH3 and CDRL3 sequence comparisons. (A) HVTN124 mAbs vs. published HIV-reactive bNAbs. The four HVTN124 mAbs are listed as rows, while the published bNAbs and their known epitopes are shown as columns. Top panel shows comparisons of the CDRH3 amino acid identity, while the bottom panel shows comparisons of the CDRL3 amino acid identity. Both scales range from 0 in white to 1.0 in black, with higher values in black corresponding to higher CDRH3 or CDRL3 sequence identity. Purple boxes signify matching V_L_ genes only, yellow boxes signify matching V_H_ genes only, and blue boxes signify matching both V genes between the respective published bNAbs and the HVTN124 mAbs. (B) HVTN124 mAbs vs. publicly available HIV-reactive antibodies. The four HVTN124 mAbs are listed on the top of each graph, with the X-axis listing CDRH3 amino acid identity and the Y-axis listing CDRL3 amino acid identity against publicly available HIV- reactive antibodies. HVTN124 mAbs were screened against 3,699 unique HIV-reactive mAb sequences (dots) from the SAbDab and PLAbDab public databases. Each colored dot relates to: grey = different V genes, purple = same V_L_ gene, yellow = same V_H_ gene, and blue = same both V genes. Blue dots were made by aligning against previously published sequences [28, 62].

Next, we compared the four HVTN124 antibodies against a compiled set of 3,699 publicly available HIV-reactive antibody sequences from the SAbDab and PLAbDab databases (Figure 7B). For each HVTN124 mAb, multiple antibodies were identified in the compiled germline dataset to contain matching light chain or matching heavy and light chains with CDRL3 amino acid identities of more than 60%, in some cases reaching 80-90%. As was the case with the high sequence signature match between 92422-98105 and DH1005, there were multiple antibodies in the compiled dataset that were encoded by both the same heavy and light chain germline genes as 92422-98105 and had high CDRH3 and CDRL3 amino acid identities (Figure 7B). Together, these results suggest that the light chain sequences for the HVTN124 mAbs are commonly found in human antibodies.

### Broadly neutralizing activity toward tier 1 and tier 2 HIV-1 strains

Next, we tested whether these antibodies neutralize HIV-1. HIV-1 neutralization was tested against a diverse panel of pseudoviruses including globally representative HIV-1 strains, additional diverse strains, and PBMC-grown SHIV viruses [55]. While three of the four antibodies did not neutralize HIV-1 pseudoviruses, antibody 92421-33628 weakly, yet broadly, neutralized HIV-1 strains belonging to tier 1 and tier 2 (Figure 8A). Interestingly, 92421-33628 also weakly neutralized murine leukemia virus (MuLV), suggesting an exceptional breadth of neutralization ability. Antibodies 92421-33628, 92422-11670, and 92422-98105 were each able to show weak neutralization when tested against tier 1 R5-tropic SHIV viruses (SHIV-1157ipEL-p) cultured in human PBMCs (hPBMCs) and/or rhesus macaque PBMCs (RhPBMC) (Figure 8A), which may represent different and more heterogeneous Env glycosylation [56].

**Figure 8.**
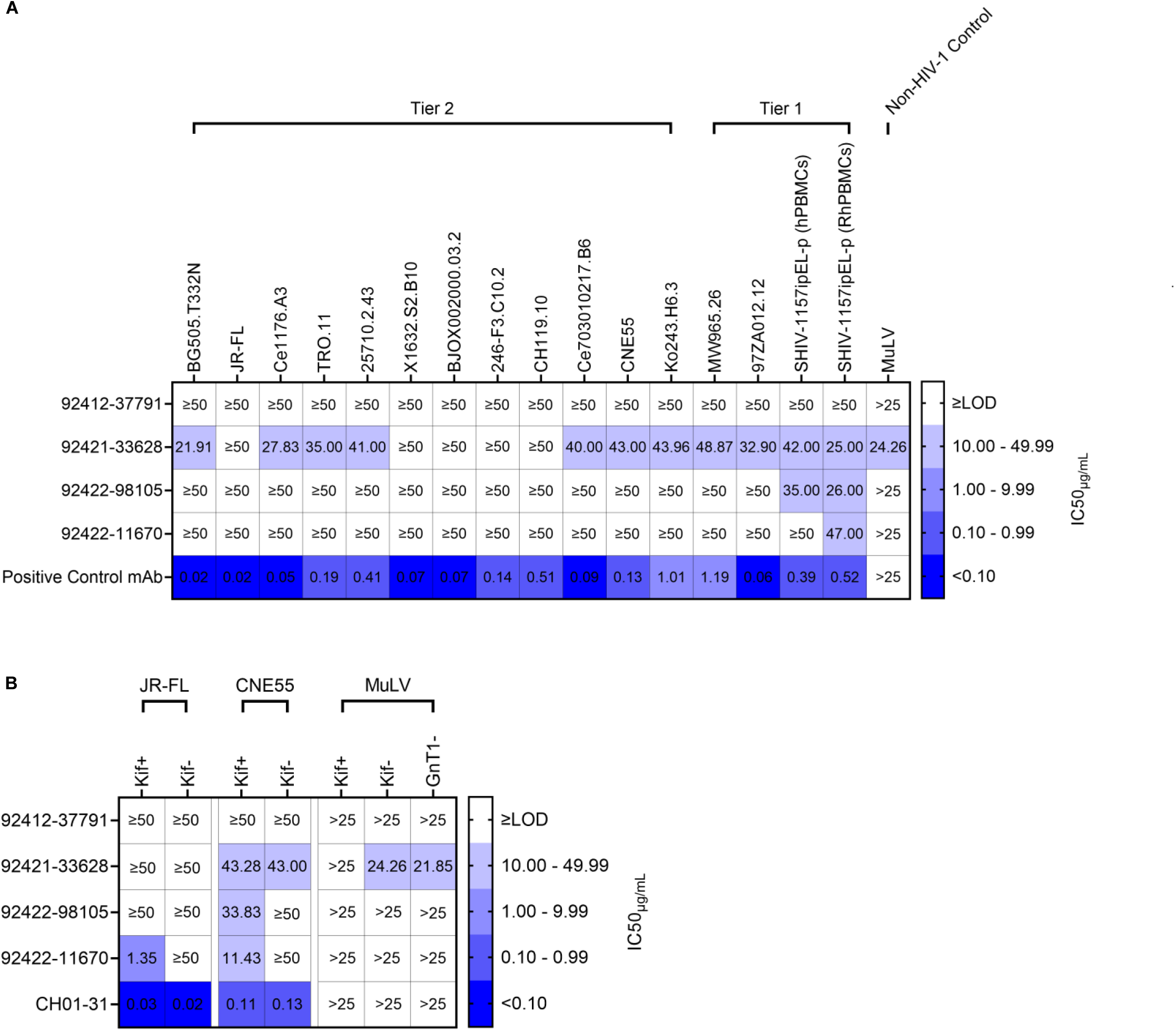
HIV-1 neutralization. (A) HVTN124 mAbs vs. HIV-1 pseudovirus and SHIV virus. HVTN124 mAbs were evaluated for neutralization against pseudotyped HIV-1 from tier 1 and tier 2 strains, which included part of a standardized global representative panel, using a TZM-bl cell neutralization assay. Murine leukemia virus (MuLV) was also included as a non-HIV-1 control. The HVTN124 mAbs are listed as rows, while each HIV-1 pseudovirus strain and MuLV is listed as columns. The positive control mAb CH01-31 was used for all conditions except for BJOX002000.03.2, which used PGT128 as the positive control mAb. For SHIV virus, VRC01 was used as the positive control mAb. IC_50_ values are listed within each cell as μg/mL mAb, with any values greater than or equal to 50 μg/mL mAb being considered as non-neutralizing against HIV-1 pseudovirus. IC_50_ values greater than 25 μg/mL mAb are considered non-neutralizing against MuLV. Heatmap key on the right is provided to categorize each IC_50_ value into either non-neutralizing in white (≥limit of detection (LOD)), weakly neutralizing in light violet (10.00 – 49.99), moderately neutralizing in dull lavender (1.00 – 9.99), potently neutralizing in dark periwinkle (0.10 - 0.99), and ultra-potently neutralizing in vivid blue (<0.10). (B) HVTN124 mAbs vs. HIV-1 pseudovirus containing modified glycans. HVTN124 mAbs were further interrogated for neutralizing activity against pseudotyped HIV-1 from tier 2 strains JR-FL and CNE55, along with MuLV, using a TZM-bl cell neutralization assay. The HVTN124 mAbs and CH01-31 positive control mAb are listed as rows, while the different HIV-1 and MuLV pseudovirus strains are listed as columns. JR-FL and CNE55 pseudoviruses were co-cultured with and without Kifunensine, which is represented as Kif+ or Kif-, respectively. MuLV was also co-cultured with and without Kifuenensine, but was additionally grown in GnT1- HEK293S mammalian cells. IC_50_ values are listed within each cell as μg/mL mAb. The heatmap key, cutoff values, and statistics are the same as in Figure 8A.

Kifunensine is a potent inhibitor of α-mannosidase I, which affects N-glycan processing to only allow cells to produce high-mannose glycans at all N-linked glycosylation sites on HIV-1 Env [57]. Because the HVTN124 mAbs target high-mannose N-glycans, we next tested their ability to neutralize kifunensine-processed HIV-1 pseudovirus. We found that three of the four HVTN124 mAbs were able to neutralize kifunensine- processed HIV-1 pseudovirus, and that kifunensine treatment improved neutralization titers in several cases (Figure 8B). Most notably, 92422-11670 was unable to neutralize JR-FL without kifunensine treatment (JR-FL Kif-), yet potently neutralized JR-FL with kifunensine treatment (JR-FL Kif+) at ∼1 µg/mL half-maximal inhibitory concentration (IC_50_). MuLV with modified glycans was also tested for neutralization, which included kifunensine-processed MuLV and MuLV grown using a N-acetylglucosaminyltransferase-deficient (GnT1-) HEK293S cell line. GnT1- cell lines arrest glycosylation at the Man5 stage, resulting in the similar production of only high-mannose glycans; however, GnT1- pseudoviruses display overall less mannose residues than when using kifunensine, giving the glycans of surface antigens of GnT1- pseudoviruses an overall reduced bulkiness [58]. Each of the three antibodies showed neutralization against either the Kif+ or GnT1- MuLV, but not both, potentially suggesting somewhat different glycan requirements for recognition by these antibodies. Full percent neutralization curves against all viruses used can be found in Figure S5A and Figure S5B. Together, these results show that HIV-1 neutralization by HVTN124 glycan-reactive antibodies can be modulated by virus glycosylation, with antibody 92421-33628 showing broad, albeit less potent, neutralization ability against a wide range of diverse HIV-1 strains, including heterologous tier 2 strains.

### Neutralizing activity against influenza, SARS-CoV-2, and Hepatitis C virus

Since the HVTN124 mAbs recognize antigens from viruses other than HIV-1, we assessed whether these antibodies could also neutralize non-HIV viruses which included: Influenza, rVSV-SARS-CoV-2 D614G, rVSV-MERS-CoV, rVSV-SARS-CoV, and HCV. None of the antibodies were found to neutralize any of the tested influenza (Figure S6) or coronavirus (Figure S7) strains. In contrast, strong neutralization activity was observed for three of the four antibodies against 14 HCV pseudoparticles (pps) from multiple genotypes across tiers 1, 2, 3, and 4 (Figure 9). These three antibodies also neutralized vesicular stomatitis virus (VSV), in agreement with the observed MuLV neutralization in the HIV-1 pseudovirus assay reported above. The breadth of the HCVpps used for neutralization included Genotype 1, Genotype 3, Genotype 4, and Genotype 5 strains to account for the broad viral diversity. Together, these results suggest that the HVTN124 glycan-reactive antibodies are capable of neutralizing diverse viruses, illuminating the extraordinary complexity of the immune system to recognize N-linked glycans on viral antigens.

**Figure 9.**
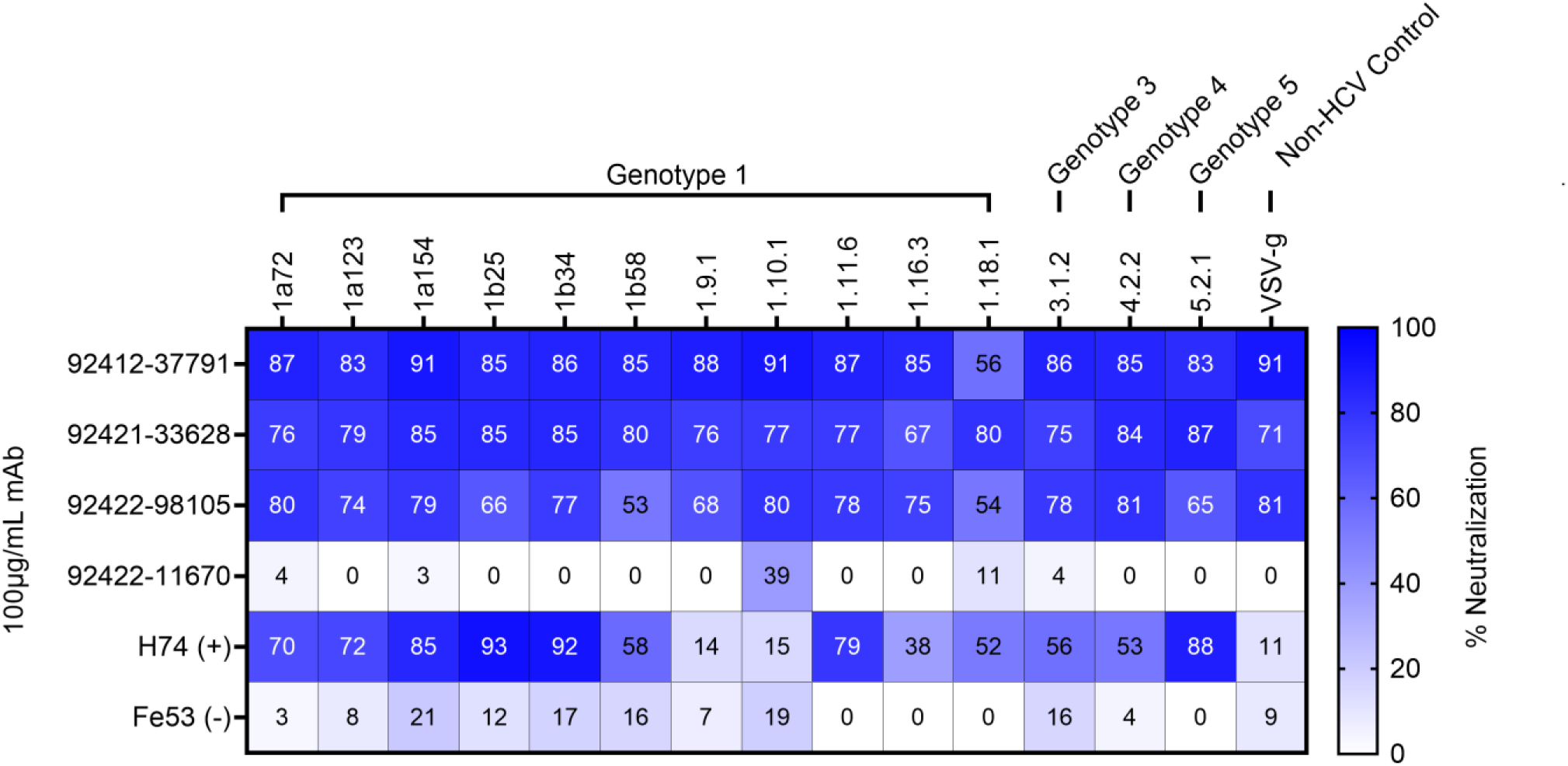
HCV neutralization. HVTN124 mAbs vs. panel of HCV pseudoparticles. HVTN124 mAbs were tested for neutralizing activity against a panel of 14 different HCVpps, along with VSV as a non- HCV control condition. The rows list each of the different mAbs used at 100 μg/mL mAb, with H74 and Fe53 being used as positive and negative controls, respectively. The columns list the different viruses, and their respective genotype, that each mAb was tested against. Viral tiers for each strain are as follows: tier 1 (5.2.1, 1.11.6), tier 2 (1a154, 1b34, 1a138, 1a123, 1.9.1, 1b25), tier 3 (1.16.3, 4.2.2, 1a72, 1.10.1, 1b58), and tier 4 (1.18.1, 3.1.2). The percent neutralization heatmap key is provided along the X- axis from 0 (white) to 100 (dark blue) percent neutralization.

### Antibody Fc effector functions against HIV-1, influenza, and SARS-CoV-2

In addition to virus neutralization, antibodies can also mediate immune responses through their Fc domain. After engaging with Fc gamma receptors (FcγRs) on effector cells, antibodies can trigger mechanisms such as antibody-dependent cellular cytotoxicity (ADCC) and/or antibody-dependent cellular phagocytosis (ADCP). Both ADCC and ADCP are two critical immune mechanisms that can aid in eliminating infected cells. The ability to elicit antibodies that can trigger ADCC and/or ADCP is of interest for HIV-1 vaccine design to provide an additional mechanism of controlling HIV- 1 infection outside of neutralization. ADCP activity of the four HVTN124 glycan-reactive antibodies was tested against HIV-1 BG505 T332N SOSIP gp140, HIV-1 CNE55 SOSIP gp140, influenza H1N1 A/New Caledonia/30/1999 HA, and SARS-CoV-2 D614G spike (Figure 10). We observed that for influenza H1N1 A/New Caledonia/30/1999 HA, antibodies 92421-33628 and 92422-11670 displayed ADCP activity above the positive threshold, albeit much weaker compared to the Fe53 and CR9114 anti-influenza control antibodies. None of the four HVTN124 mAbs displayed detectable ADCP activity against SARS-CoV-2 D614G spike. When tested against HIV-1 BG505 N332T SOSIP gp140, ADCP activity was observed for three of the four antibodies, albeit again weaker than other known HIV-1 antibodies.

**Figure 10.**
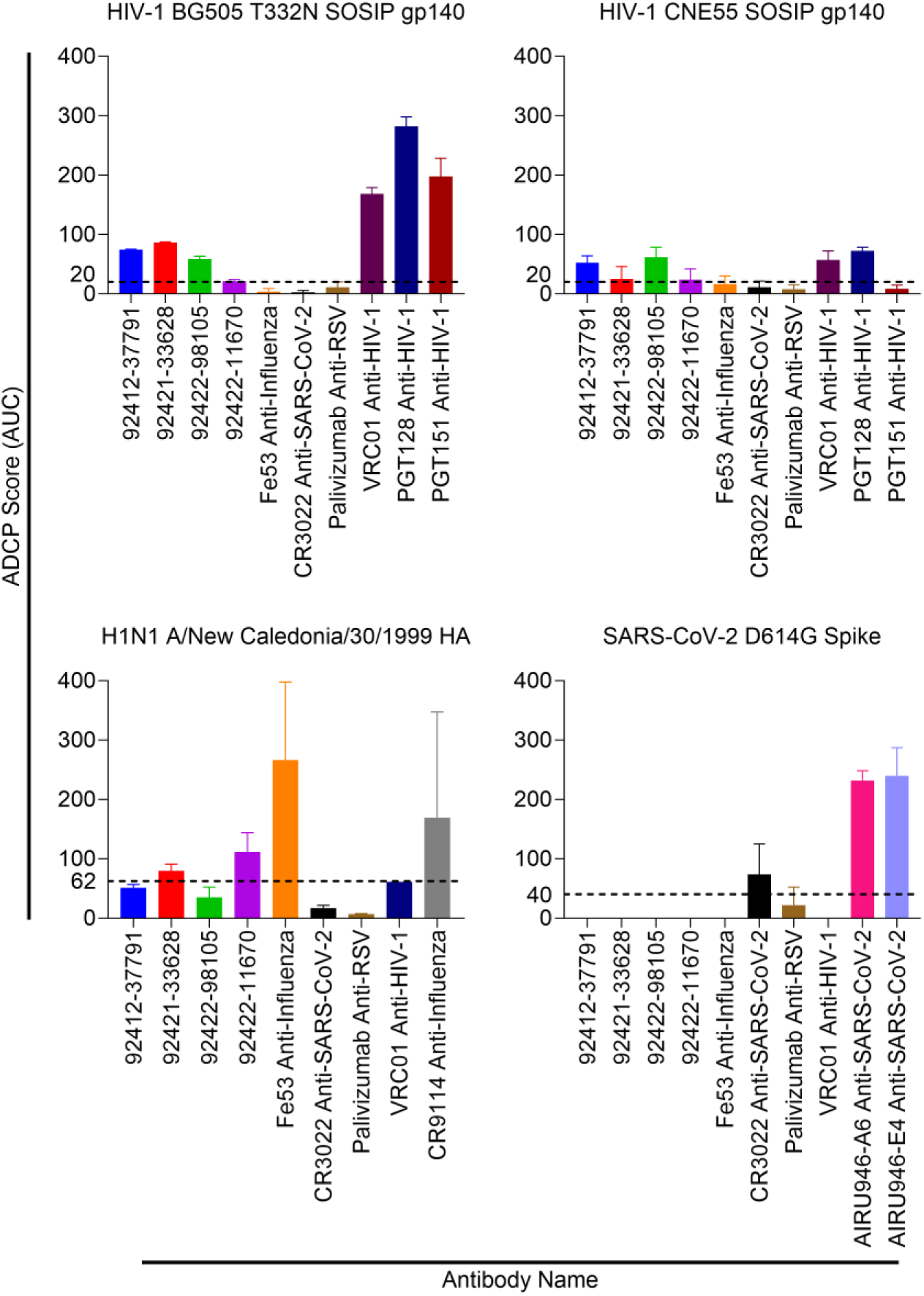
HVTN124 mAb ADCP activity. ADCP activity was measured against HIV-1 BG505 T332N SOSIP gp140, HIV-1 CNE55 SOSIP gp140, influenza H1N1 A/New Caledonia/30/1999 HA, and SARS-CoV-2 D614G Spike. Antigens were each biotinylated and conjugated to fluorescent beads with immobilized streptavidin, then mixed with each mAb. Antibodies are listed along the X- axis, while the Y-axis lists the ADCP score as AUC. The control mAbs used include: VRC01, PGT128, PGT151 = HIV-1; Fe53, CR9114 = influenza; CR3022, AIRU946-A6, AIRU946-E4 = SARS-CoV-2. Palivizumab was included as a negative control antibody. THP-1 monocytes were subsequently added and ADCP was calculated based on the engulfment of the antigen-coated beads by the THP-1 cells as measured by flow cytometry. Antigen engulfment is indicated as an ADCP score.

ADCC activity of the four HVTN124 glycan-reactive antibodies was tested against HIV-1 BG505 T332N SOSIP gp140, influenza H1N1 A/New Caledonia/30/1999 HA, and SARS-CoV-2 D614G spike (Figure 11). We observed that for all three antigens, antibodies 92412-37791 and 92422-11670 displayed the strongest ADCC activity above the positive threshold. Antibody 92422-98105 also displayed ADCC activity above this threshold against all three antigens, but was much weaker than 92412-37791 and 92422-11670. Together, these results show that all four HVTN124 glycan-reactive antibodies can mediate Fc effector functions, albeit with differing strengths in ADCP and ADCC responses.

**Figure 11.**
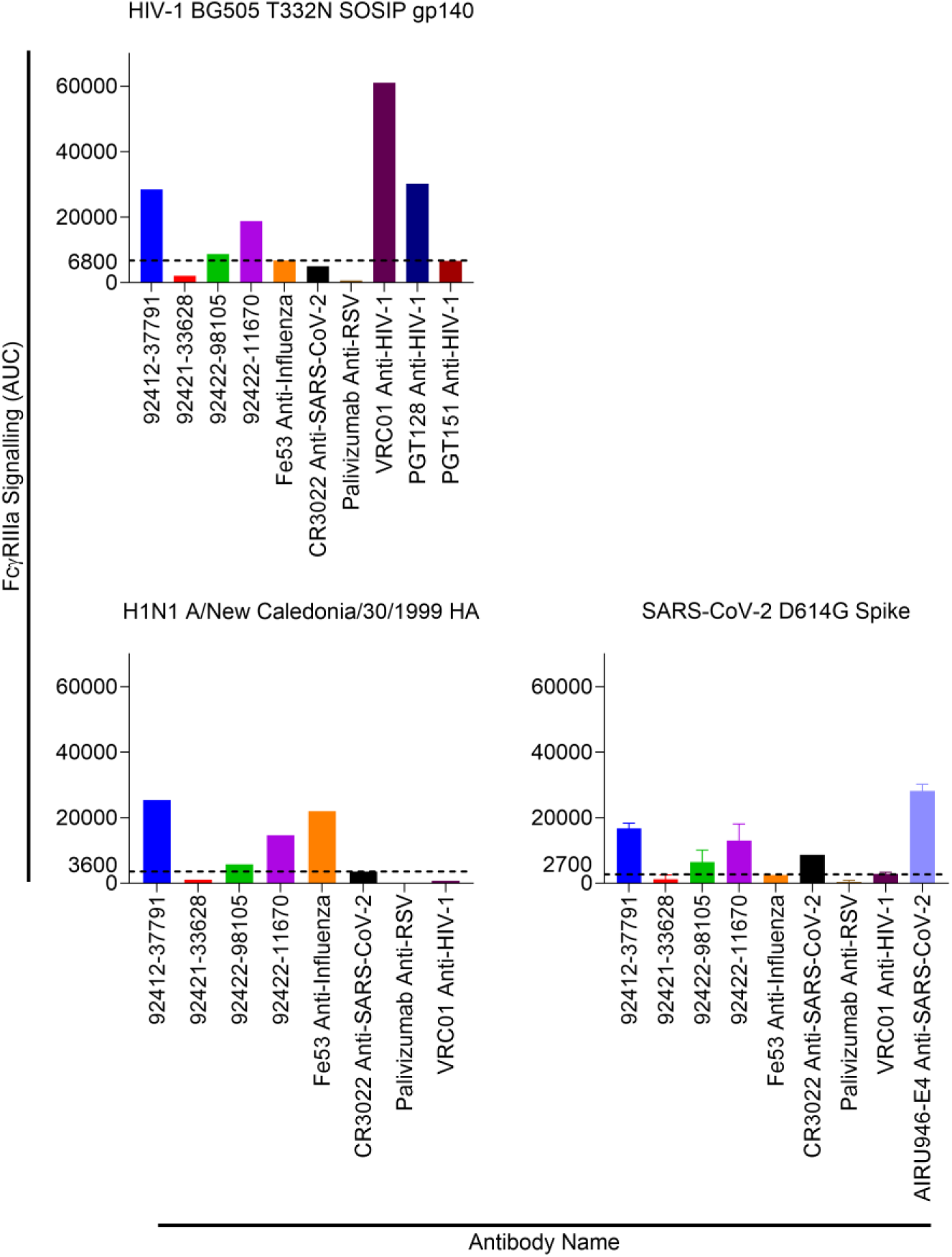
HVTN124 mAb ADCC activity. ADCC activity was measured against HIV-1 BG505 T332N SOSIP gp140, influenza H1N1 A/New Caledonia/30/1999 HA, and SARS-CoV-2 D614G Spike. Antibodies are listed along the X-axis, while the Y-axis lists the FcγRIIa signaling as AUC. The control mAbs used include: VRC01, PGT128, PGT151 = HIV-1; Fe53 = influenza; CR3022, AIRU946-E4 = SARS-CoV-2. Palivizumab was included as a negative control antibody. Fc crosslinking activity is reported as AUC.

## Discussion

Antibody responses to HIV-1 infection have been extensively studied, resulting in the identification of a wide range of broadly neutralizing antibodies, new insights into the genetic and structural features associated with effective protection against infection, the discovery of new sites of virus vulnerability as potential templates for antibody-based vaccines. In contrast, there is still limited understanding of antibody responses to HIV-1 vaccination in humans. Similarly, mapping the role of pre-existing antibody repertoires on modulating responses to HIV-1 vaccination is also not clear, with extensive focus to date on identifying rare precursors for difficult to elicit broadly neutralizing antibodies [59]. Yet, the antibody repertoire in HIV-naive humans is highly complex, comprising antibodies with diverse sequence features and phenotypes that have the potential to engage with the vaccine antigens upon immunization.

Recent work suggests that one such class of antibodies that have the capability of engaging HIV-1 vaccine antigens in HIV-naïve individuals are glycan-reactive FDG antibodies [28]. Multiple efforts have previously described how glycan-reactive antibodies uniquely display broad reactivity across multiple viral families [28, 60, 61]. As such, FDGs (and, more generally, glycan-reactive) antibodies may represent an immune system mechanism that provides a first line of defense against a broad range of viral pathogens in humans. At the same time, this class of antibodies may also be among the first to engage with a vaccine antigen upon immunization, highlighting the importance of deciphering the features and functional phenotypes associated with these antibodies.

In this report, we discovered and characterized a set of glycan-reactive antibodies isolated from participants in the HVTN124 HIV-1 vaccine trial. While these antibodies exhibited stronger binding to the trimeric Env-derived SOSIP antigens, binding to the gp120 HVTN124 vaccine antigens was also observed. Notably, for three of the four antibodies, reactivity against the HIV-1 antigens was not observed when reverted to germline. This potentially suggests that these antibodies could have been part of the pre-existing repertoire that could engage with the vaccine, rather than being elicited by the vaccine itself. Although, other factors such as BCR vs. monoclonal antibody affinity and potential differences in true vs. inferred germline precursor sequences could also play a role, preventing any definitive causal conclusions about the generation of these antibodies.

Among the four isolated antibodies, 92421-33628 appeared to have the most intriguing and unique profile. This antibody showed an ability to neutralize, albeit weakly, a diverse set of HIV-1 viruses from tiers 1, 2, and 3 when tested in a pseudovirus assay, while not showing signs of autoreactivity. To our knowledge, this is the first report of a glycan-reactive broadly neutralizing antibody that was not obtained from an HIV infection sample, demonstrating that these types of antibodies can be elicited outside of an HIV-1 infection. Interestingly, and in line with prior studies, three of the antibodies showed neutralization ability against kifunensine-treated pseudovirus, suggesting that the type of glycans present on the surface of the virus can modulate neutralization by these types of glycan-reactive antibodies [28]. Importantly, these three antibodies also exhibited neutralization of SHIV viruses cultured from PBMCs, which represent different and potentially more heterogeneous glycosylation compared to HIV-1 pseudovirus produced in cell lines. Further, one of these neutralizing antibodies (92422-98105) showed high sequence similarity to a previously published glycan-reactive antibody in both the heavy and light chain sequences, as part of a public class of glycan HIV-reactive antibodies.

Together, the results reported here build on current knowledge about the features and functional phenotypes of glycan-reactive antibodies and their potential role as a component of effectively responding to HIV-1 vaccination in HIV-naïve repertoires. These efforts motivate further research toward mechanistically understanding the molecular and developmental basis of this glycan-reactive antibodies, along with how this class of antibodies can be better targeted for large-scale engagement in future HIV- 1 vaccine trials.

## Methods

### Immunogen panel expression, purification, and validation

HIV-1 immunogens were designed using the SOSIP platform to yield soluble Env proteins that are stabilized in the pre-fusion conformation. SOSIP proteins include the following mutations: an intermolecular disulfide bond between gp120 and gp41 (A501C and T605C), a trimer-stabilizing mutation (I559P), a truncated gp41 transmembrane region at position 664, an I201C/A433C mutation to inhibit CD4-induced movement of Env, and a variable flexible serine-glycine linker between gp120 and gp41 (positions 507 and 512) to create single-chain constructs [30]. SOSIP immunogens were transiently transfected into Expi293F cells in FreeStyle F17 expression media (Thermo Fisher). Supplements were added at a final concentration of 0.1% Pluronic Acid F-68 and 1X Glutamax. Cells were transfected using the Expifectamine transfection reagent (Thermo Fisher Scientific), followed by culturing in a shaker for 5 to 7 days at 8% CO_2_ at 37°C. Following transfection, cell cultures were centrifuged at 4,000 x *g* for 15 minutes to form a stable cell pellet. Centrifuged cell culture supernatant was then filtered using Nalgene Rapid Flow Disposable Filter Units with PES membrane (0.45µM), then run slowly over a chromatography column with agarose-bound Galanthus nivalis lectin (Vector Laboratories cat no. AL-1243-5) at 4°C. After the supernatant had flowed through, the column was washed with 1x PBS, then the immunogen was eluted with 30 mL of 1 M methyl α-D-mannopyranoside. The eluted immunogen was then buffer exchanged in 1x PBS three times, followed by concentrating using 100 kDa Amicon Ultra centrifugal filter units. The concentrated immunogen was run through a Superdex 200 Increase 10/300 GL on the AKTA FPLC system. Any peaks indicative of Env trimers were collected based on elution volume, followed by validation of molecular weight using SDS-PAGE gel electrophoresis and antigenicity using an ELISA format against Env-specific antibodies.

### Antigen biotinylation

All immunogens used for LIBRA-seq were non-specifically biotinylated using the EZ-Link Sulfo-NHS-Biotinylation Kit (Thermo Scientific cat no. 21425) according to the manufacturer’s protocol.

### Oligonucleotide barcodes and conjugation to antigens

Oligonucleotides used in the LIBRA-seq pipeline contain a 15 base pair barcode that is unique for each antigen in the panel. During sample processing with the 10X Genomics platform, the part of the oligonucleotide sequence will anneal to 10X Genomics bead, specifically at the template switch oligonucleotide. Lastly, each oligonucleotide contains a truncated TruSeq small RNA sequence as the following: 5′-CCTTGGCACCCGAGAATTCCA<u>NNNNNNNNNNNNNNN</u>CCCATATAAGA*A*A-3′ N’s in this example represent the unique barcode for distinguishing each antigen. Oligonucleotides were ordered from Sigma-Aldrich and IDT with a 5’ amino modification and were HPLC purified. During LIBRA-seq analysis, the following unique barcodes were used with their respective antigens: HVTN124 Clade A gp120 (GCTCCTTTACACGTA), BG505 N332T SOSIP gp140 (CAGATGATCCACCAT), KNH1144 SOSIP gp140 (ACAATTTGTCTGCGA), HVTN124 Clade B gp120 (AGACTAATAGCTGAC), B41 SOSIP gp140 (TCACAGTTCCTTGGA), TRO.11 SOSIP gp140 (AACCTTCCGTCTAAG), HVTN124 Clade C gp120 (TACGCCTATAACTTG), CZA97 N332T SOSIP gp140 (TAACTCAGGGCCTAT), ZM106.9 SOSIP gp140 (CAGCCCACTGCAATA), HVTN124 Clade AE gp120 (CTTCACTCTGTCAGG), CNE55 SOSIP gp140 (TCCTTTCCTGATAGG), influenza H1N1 A/New Caledonia/30/1999 HA (AATCACGGTCCTTGT)

Each unique oligonucleotide barcode was directly conjugated to their respective antigen using the Solulink Protein-Oligonucleotide Conjugation kit (TriLink, S-9011) according to the manufacturer’s protocol. The antigen and oligonucleotide were first desalted, then the oligonucleotide was modified with a 4FB crosslinker while the biotinylated antigen was separately modified with S-HyNic. These two products were then mixed and incubated to allow a stable bond to form between the antigen and oligonucleotide. Concentrations of the antigen-oligonucleotide conjugates were determined using a BCA assay, followed by measuring the S-HyNic and 4FB molar substitution ratios of each conjugate using the NanoDrop based on the Solulink protocol. AKTA FPLC was then used to remove excess oligonucleotide from the antigen-oligonucleotide conjugates to reduce noise during Illumina Next-Generation Sequencing by VANTAGE, with the removal of excess oligonucleotide being validated using SDS-PAGE silver stains.

### 10X Genomics single cell processing for Illumina next-generation sequencing

Following single-cell sorting, samples were loaded onto a Chromium Controller microfluidics device (10X Genomics) and processed as previously described above. A target capture of 10,000 B cells was used based on the recommended specifications for the 10X Genomics platform.

### Bioinformatics analysis

By utilizing our lab’s established R studio pipeline, the paired-end FASTQ files from the oligonucleotide libraries were input to first process and label cell barcode reads. This initial step results in a matrix of unique molecular identifiers (UMIs) and antigen barcodes. Next, files related to B cell receptor contigs were processed using CellRanger 3.1.0 (10X Genomics) and GRCh38 V(D)J 7.0.0 as a reference, with the antigen barcode libraries processed in the earlier step also being processed using CellRanger (10X Genomics). Any cell barcodes that overlapped with the antigen barcodes and UMIs were then subject to further analysis. Cell barcodes were eliminated from the matrix if they contained non-functional heavy and/or light chain sequences, in addition to if the cell barcodes were associated with multiple functional heavy and/or light chain sequences, with both criteria aiming to eliminate any multiplets. B cell receptor contigs (file output by CellRanger as filtered_contigs.fasta) were aligned to IMGT reference genes using HighV-Quest. This output was analyzed using ChangeO, then combined with the antigen barcode UMI score matrix. The LIBRA-seq score (LSS) for each antigen in the panel against every B cell was then calculated as previously described [29].

### Enzyme linked immunosorbent assay (ELISA)

ELISA for immunogens was done in a Immulon® 2 HB 96-Well Microtiter EIA Plate. Immunogens were diluted to 2 µg/mL in 1% BSA in PBS-T, then added to the plates and incubated overnight at 4°C. The next day, plates were washed using 0.05% Tween20 (PBS-T) to remove any unbound antigen, then blocked in 5% BSA in PBS-T for two hours at room temperature. After washing again after incubating, antibody dilution series in 1% BSA in PBS-T were plated and incubated for one hour at room temperature, then washed. The secondary antibody, goat anti-human IgG conjugated to peroxidase, was applied to each well at a 1:10,000 dilution in 1% BSA in PBS-T, then incubated for one hour at room temperature. After washing again, plates were developed by adding TMB substrate to each well for five minutes at room temperature. The reaction was quenched by adding 1N sulfuric acid, where plates were then read at 450nm. Values were then subtracted from the average A450nm values from the no-mAb condition to account for background noise. Concentrations were then transformed, where µg/mL mAb was converted to ng/mL mAb, which the log10 of ng/mL mAb was taken to give a new concentration as log[mAb] (ng/mL). Area under the curve was then taken, where the baseline was set to Y = 0, minimum peak height was set to ignore peaks that are less than 10% of the distance from minimum to maximum Y, and peak direction was defined as having peaks that went above baseline.

### Mannose-competition ELISA

Mannose competition ELISAs were performed as described above, but with minor modifications. Following coating with CNE55, plates were blocked for 1 hour at room temperature in either 1M D-(+)-mannose 5% BSA PBS-T or 5% BSA PBS-T. Primary antibodies were prepared in either 1M D-(+)-mannose 1% BSA PBS-T or 1% BSA PBS- T, starting at 10 µg/mL with a 5-fold serial dilution. After washing three times, primary antibodies were added to each plate for 1 hour at room temperature. Following another wash step, goat anti-human IgG conjugated to peroxidase was added at a 1:10,000 dilution in 1% BSA PBS-T for 1 hour at room temperature. After washing again, plates were developed and quenched as described above. Values were also transformed as described previously.

### Enzymatic deglycosylation of HIV-1 CNE55 SOSIP gp140

The glycan-dependency of antibodies was evaluated by treating antigen under different condition, then testing by ELISA. Antigen under the native condition were quickly thawed after being taken from –80°C storage and had no enzymatic treatment applied. The 37°C condition had antigen diluted in 1X Glycobuffer 2, then incubated at 37°C overnight. The 37°C + PNGase F condition had antigen diluted in 1X Glycobuffer 2 and incubated with PNGase F at 37°C overnight. The 37°C + Endo H condition had antigen diluted in 1X Glycobuffer 2 and incubated with Endo H at 37°C overnight. The 37°C + O Glycosidase condition had antigen diluted in 1X Glycobuffer 2 and incubated with O Glycosidase at 37°C overnight. The 95°C condition had antigen diluted in Glycoprotein Denaturing Buffer and boiled at 95°C for 10 minutes, then chilled on ice for 2 minutes. 1 µL of enzyme was used per 10 µg of total antigen for each condition. The different antigen conditions were then directly coated onto ELISA plates and assayed as previously described in the ELISA methods section. Values were also transformed as described previously. To normalize values against the native condition, values gathered from calculating the AUC were applied to the normalization analysis option in GraphPad Prism. Settings used included setting 0% defined as Y = 0, 100% defined as the AUC value from the native condition for each separate antibody, and presenting the results as percentages. This resulted in the native condition being set at 100% for all antibodies across the different conditions.

### AtheNA autoantigen panel

Monoclonal antibody reactivity was measured using the AtheNA Multi-LyteANA- IIPlustestkit (Zeusscientific,Inc.), which tested against nine autoantigens (SSA/Ro, SS- B/La, Sm, ribonucleoprotein (RNP), Scl 70, Jo-1, dsDNA, centromereB, and histone). Antibodies were incubated with AtheNA beads for 30 minutes at concentrations of 50, 25, 12.5, or 6.25 µg/mL mAb. After washing beads, they were incubated with secondary and read on the Luminex platform as specified in the kit protocol. AtheNA software was used to analyze the data. Positive samples were those that received a score >120, while negative samples received a score of <100. Values between 100 and 120 were considered intermediate.

### Cardiolipin reactivity

Monoclonal antibody reactivity to cardiolipin was measured using the Quanta Lite ACA IgG III kit (Inova Diagnostics, Inc.) by diluting antibodies to final concentrations of 100, 50, 25 and 12.5 µg/mL in sample diluent. 100 µL of each sample was added to assay plates along with kit controls and the kit SOP was followed for the remainder of the assay. Phospholipid units (GPL) were calculated against a linear standard curve per kit instructions. An individual well was positive if GPL was >20, negative if <15 and indeterminate if between 15 to 20. Any given antibody needed to be positive for two consecutive wells (*i.e.*, to at least 50 µg/mL) to be considered cardiolipin positive.

### HEp-2 cell binding

Monoclonal antibody binding to whole, un-permeabilized, un-infected HEp-2 cell monolayer cultures was performed via flow cytometry. HEp-2 cells were collected and washed three times with DPBS-BSA before counting. Roughly 1 million cells/condition were stained to a final concentration of 100, 10, or 1 µg/mL mAb diluted in DPBS-BSA for 20 minutes at 4°C. Cells were washed three times with DPBS-BSA, then stained with anti-IgG FITC (Southern Biotech) diluted at 1:1,000 in DPBS-BSA, along with DAPI to stain the cell nuclei, for 20 minutes at 4°C. Cells were washed and fluorescence was acquired on a 4-Laser Fortessa (BD Biosciences). FCS files were analyzed, and figures were generated using CytoBank.

### NSEM

Antibodies from -80°C were separately thawed at RT in an Al block for 5 min. Samples were then diluted to 20 µg/mL with 0.02 g/dL Ruthenium Red in HBS (20 mM HEPES, 150 mM NaCl pH 7.4) buffer. After 10 to 15 minute incubation, samples were applied to a glow-discharged carbon-coated EM grid for 8-10 second, blotted, consecutively rinsed with 2 drops of 1/20x HBS, and stained with 2 g/dL uranyl formate for 1 minute, blotted and air-dried. Grids were examined on a Philips EM420 electron microscope operating at 120 kV and nominal magnification of 49,000×, and 20 images were collected on a 76 Mpix CCD camera at 2.4 Å/pixel. Images were analyzed by 2D class averages using standard protocols with Relion 3.0 [63].

### Published and public antibody analysis

The published and public reference libraries use paired heavy chain and light chain human BCR sequences from published sources, and the SAbDab and PLAbDab public databases, respectively. We used custom Python scripts utilizing NumPy and Pandas to determine published and public clones between our data and a reference library. The identification of published and public clonotype similarity utilized a cutoff of 70% amino acid sequence identity for the CDRH3 and CDRL3 regions, in addition to containing matching heavy variable (V_H_), light variable (V_L_), and joining (J) gene usage. Levenshtein distance was used for comparison in CDR3 identities. When comparing CDRH3 and CDRL3 of different lengths, gaps are penalized the same as mismatches.

### HIV-1 and MuLV pseudovirus and SHIV virus production and neutralization assays

HIV-1 and MuLV pseudoviruses were produced by co-transfecting HEK293T cells or GlcNAc transferase I enzyme-deficient 293S cells (293S/GnTI-) with an Env-expressing plasmid and an Env-deficient HIV-1 backbone vector pSG3ΔEnv (BEI #11051) as previously reported [64]. SHIV virus (SHIV-1157ipEL-p) was cultured using CD8- depleted human or rhesus PBMCs as previously reported [56] The optimal tissue culture infectious dose (TCID) of each pseudovirus or SHIV was determined by titrating in TZM-bl cells. Antibody neutralization was further characterized using the TZM-bl cell-based assay [65]. This standardized assay quantifies the inhibition of TZM-bl cell infection by Env-pseudotyped viruses through antibody-mediated activity. The panel of viruses included tier 1 and tier 2 strains, a murine leukemia virus (MuLV), and PBMC-grown SHIV viruses. Virus neutralization was measured as a function of reductions in luciferase (Luc) reporter gene expression after a single round of infection in TZM-bl cells. Briefly, a pre-titrated dose of virus was incubated with serial diluted antibodies in duplicate for 1 hour at 37°C in 96-well plates, followed by addition of freshly trypsinized TZM-bl cells containing DEAE-Dextran. Assay plates are incubated at 37°C for 44 to 72 hours, allowing a single round of infection in TZM-bl cells. Results are displayed as the antibody concentration at which relative luminescence units (RLU) is reduced to 50% (IC_50_) compared to virus control wells.

### H1N1 and H3N2 Influenza virus production and neutralization assay

Production of replication-restricted reporter (R3) H1N1 and H3N2 Influenza viruses was described previously [66]. Viral genomic RNA encoding functional PB1 (R3ΔPB1 viruses) was replaced with a gene encoding a fluorescent protein. R3ΔPB1 viruses were rescued and propagated in cell lines stably expressing PB1 via reverse genetics. Viral stocks were titered by determining the number of fluorescent units per mL (FU/mL). Each mAb was diluted in Opti-MEM to 100 μg/mL, serially diluted, mixed 1:1 with viruses, and allowed to co-incubate for 1 hour in a humidified 37°C incubator (5% CO_2_). Viruses were diluted to 1x10^4^ to 4x10^4^ FU/mL in Opti-MEM prior to mixing. MDCK-SIAT1 cells (Millipore Sigma #05071502) expressing PB1 were then harvested, washed in Opti-MEM, diluted to 1x10^6^ cells/mL, added to the mAb-virus mixtures (3:1 mAb-virus to cells), then immediately transferred to a 384-well plate format in quadruplicate. Plates were then incubated for 18 to 22 hours at 37°C in a humidified incubator (5% CO_2_). After incubation, the fluorescent units of each well were counted using a Celigo Image Cytometer (Nexcelom) with a customized red filter to detect mKate2/TdKatushka2 reporter signals. The percent neutralization was calculated by constraining the virus control (virus + cells) as 0% and the cell control (cells only) as 100% and plotted against mAb concentration.

### rVSV-SARS-CoV-2, rVSV-MERS-CoV, and rVSV-SARS-CoV virus production

The generation of a replication-competent VSV expressing S proteins of respective CoVs that replaces the VSV G protein was described previously [67]. The S protein-expressing VSV viruses were propagated in either Vero-E6 or MA104 cell culture monolayers (African green monkey, ATCC CRL-2378.1) as described previously, and viral stocks were titrated on Vero E6 cell monolayer cultures [67]. VSV plaques were visualized using neutral red staining.

### rVSV-SARS-CoV-2, rVSV-MERS-CoV, and rVSV-SARS-CoV neutralization using Real-time cell analysis (RTCA)

To determine the neutralizing activity of IgGs, we used real-time cell analysis (RTCA) assay on an xCELLigence RTCA MP Analyzer (ACEA Biosciences Inc.) that measures virus-induced cytopathic effect (CPE) as previously described [68]. Briefly, 50 μL of cell culture medium (DMEM supplemented with 2% FBS) was added to each well of a 96- well E-plate using a ViaFlo 384 liquid handler (Integra Biosciences) to obtain background reading. A suspension of 18,000 Vero-E6 cells in 50 μL of cell culture medium was seeded in each well, and the plate was placed on the analyzer. Measurements were taken automatically every 15 minutes, and the sensograms were visualized using RTCA software version 2.1.0 (ACEA Biosciences Inc). Respective VSV-CoV (0.05 MOI, 150 PFU per well) was mixed 1:1 with a dilution of mAb in a total volume of 100 μL using DMEM supplemented with 2% FBS and incubated for 1 hour at 37°C in 5% CO2. At 16 to 18 hours after seeding the cells, the virus-mAb mixtures were added in triplicates to the cells in 96-well E-plates. Triplicate wells containing virus only (for maximal CPE in the absence of mAb) and wells containing only Vero cells in medium (for no-CPE wells) were included as controls.

Plates were measured continuously (every 15 minutes) for 48 to 72 hours to assess virus neutralization. Normalized cell index (CI) values at the endpoint (48 to 72 hours after incubation with the virus) were determined using the RTCA software version 2.1.0 (ACEA Biosciences Inc.). Results were expressed as percent relative infection in the presence of respective mAb, relative to control wells with no CPE minus CI values from control wells with maximum CPE. IC50 values were determined by nonlinear regression analysis using Prism software.

### HCV pseudoparticle (HCV_PP_) production and neutralization assay

A panel of 14 HCVpps from genotype 1 (UKNP1.11.6, UKNP 1A154, UKNP1B34, UKNP1.9.1, UKNP1A123, UKNP1B25, UKNP1.16.3, UKNP1A72, UKNP1.10.1, UKNP1B58, UKNP1.18.1), genotype 3 (UKNP3.1.2), genotype 4 (UKNP4.2.2), and genotype 5 (UKNP5.2.1) from were produced via lipofectamine-mediated transfection of pNL4-3.Luc.R-E, HCV E1E2, and pAdVantage plasmids into HEK293T cells [69, 70]. Vesicular stomatitis virus (VSV) was included as an HCV-specific control. For testing neutralization, 96-well solid flat white bottom polystyrene TC-treated microplates had 8,000 Hep3B cells applied per well, then incubated overnight at 37°C. To test each mAb, HCVpps were co-incubated with 100 µg/mL mAb for 1 hour, then added in duplicate to Hep3B target cells for 5 hours. Following incubation, medium was changed to 100µL of phenol-free Hep3B media, then incubated for 72 hours at 37°C. Infectivity was later quantified using a luciferase-based assay measured in relative light units (RLUs) in Berthold Luminometer (Berthold Technologies Centro LB960). Percent neutralization for each mAb was calculated as [1-(RLUmAb/ RLUIgG)] x 100]. Percent neutralization for the dilution curves and was calculated as [1-(RLUmAb/RLUPBS)] x 100. H74 and Fe53 were included as positive and negative controls, respectively.

### Antibody-dependent cellular phagocytosis (ADCP) assay

The BirA biotin-protein ligase bulk lyophilized reaction kit was used to biotinylate avi-tagged proteins to ensure correct orientation of the antigen and coated on fluorescent neutravidin beads as previously described (Ackerman, et al., 2011). Antigen coated beads were incubated for two hours at 37°C, 5% CO_2_ with a titration of monoclonal antibodies at 2 μg/mL and titrated five-fold. Beads were washed and further incubated with THP-1 cells overnight, fixed with Paraformaldehyde (PFA) (Sigma) and interrogated on the Cytoflex (Beckman Coulter). ADCP score was calculated as a proportion of THP- 1 cells that internalized beads multiplied by the geometric mean fluorescence intensity of the population. Signal from a “no antibody” negative control was subtracted to remove background signal.

### Antibody-dependent cellular cytotoxicity (ADCC) assay

Antibodies were screened for their capacity to cross-link and activate the FcγRIIIa (CD16) receptor on the surface of Jurkat cells and D614G SARS-CoV-2 spike expressing cells or BG505 or NC/99 trimer coated on to plates as a proxy for ADCC. HEK293T cells were transfected with 5 μg of SARS-CoV-2 ancestral variant spike (D614G) using 1 mg/mL PEI-MAX 40,000 (Polysciences) and incubated for 2 days at 37°C. Alternatively BG505 protein was coated at 1 μg/mL on a high binding ELISA 96- well plate and incubated at 4 °C overnight. Plates were then washed with PBS and blocked at room temperature for 1 hour with PBS + 2.5% BSA. Subsequently, protein or 1 × 10^5^ spike-transfected cells per well were incubated with monoclonal antibodies (in RPMI 1640 medium supplemented with 10% FBS 1% Pen/Strep (Gibco, Gaithersburg, MD) for 1 hour at 37 °C. To confirm spike expression from the HEK293T cells, binding of CR3022 and 946-A6 was detected by anti-IgG APC staining and measured by flow cytometry. Twenty µl of supernatant was transferred to a white 96-well plate with 50 μL of reconstituted QUANTI-Luc secreted luciferase and read immediately on a Victor 3 luminometer with 1s integration time. Additionally, 1x cell stimulation cocktail (Thermo Fisher Scientific, Oslo, Norway) and 2 μg/mL ionomycin in R10 were added as controls to induce the transgene and confirm sufficient expression of the Fc receptor.

### Quantification and statistical analysis

ELISA (standard error of the mean; SEM), ADCP/ADCC, and neutralization error bars shown were calculated using GraphPad Prism version 10.4.0.

## Acknowledgements

We gratefully thank the members of the Georgiev lab for their invaluable assistance and constructive comments. Special thanks to David Flaherty, Olivia Murfield, Emma McLaughlin, and Brittany Matlock from the VUMC Flow Cytometry Shared Resource for their assistance with cell sorting. The VUMC Flow Cytometry Shared Resource is supported by the Vanderbilt Ingram Cancer Center (P30 CA68485) and the Vanderbilt Digestive Disease Research Center (DK058404). We also appreciate Angela Jones, Jamie Roberson, Latha Raju, and Lana Olson from the Vanderbilt Technologies for Advanced Genomics Core (VANTAGE) for providing technical expertise in library construction and sequencing. VANTAGE is supported by grants from organizations including CTSA (5UL1 RR024975-03), Vanderbilt-Ingram Cancer Institute (P30 CA68485), Vanderbilt Vision Center (P30 EY08126), and NIH/NCRR (G20 RR030956). We thank Shan Lu and the HVTN124 program team for providing PBMCs used in this study. In addition, we acknowledge the financial support of the National Institutes of Health (NIH) R01 AI152693-04, allowing the findings described in this manuscript to be possible. The funds allocated for this research played no role in the conceptualization or execution of any studies, or regarding manuscript preparation.

## Author Contributions

Conceptualization and Methodology: P.J.J. and I.S.G.; Investigation: P.J.J., X.S., A.A.A., P.T.W., K.J., R.J.E., G.S., S.I.R., N.P.M., S.L., M.B., R.A.G., J.M., N.S., T.A.S., S.Z., R.P., S.F., A.K.J., B.N.H., Y.P.S.; Writing – Original Draft: P.J.J. and I.S.G.; Writing – Review and Editing: All authors; Funding Acquisition: P.J.J. and I.S.G.; Resources: R.M.R., J.E.C., R.H.C., J.B., K.M., B.F.H., P.L.M, P.A., D.C.M., S.A.K., S.L., I.S.G.; Supervision: P.J.J. and I.S.G.

---

**Figure S1.**
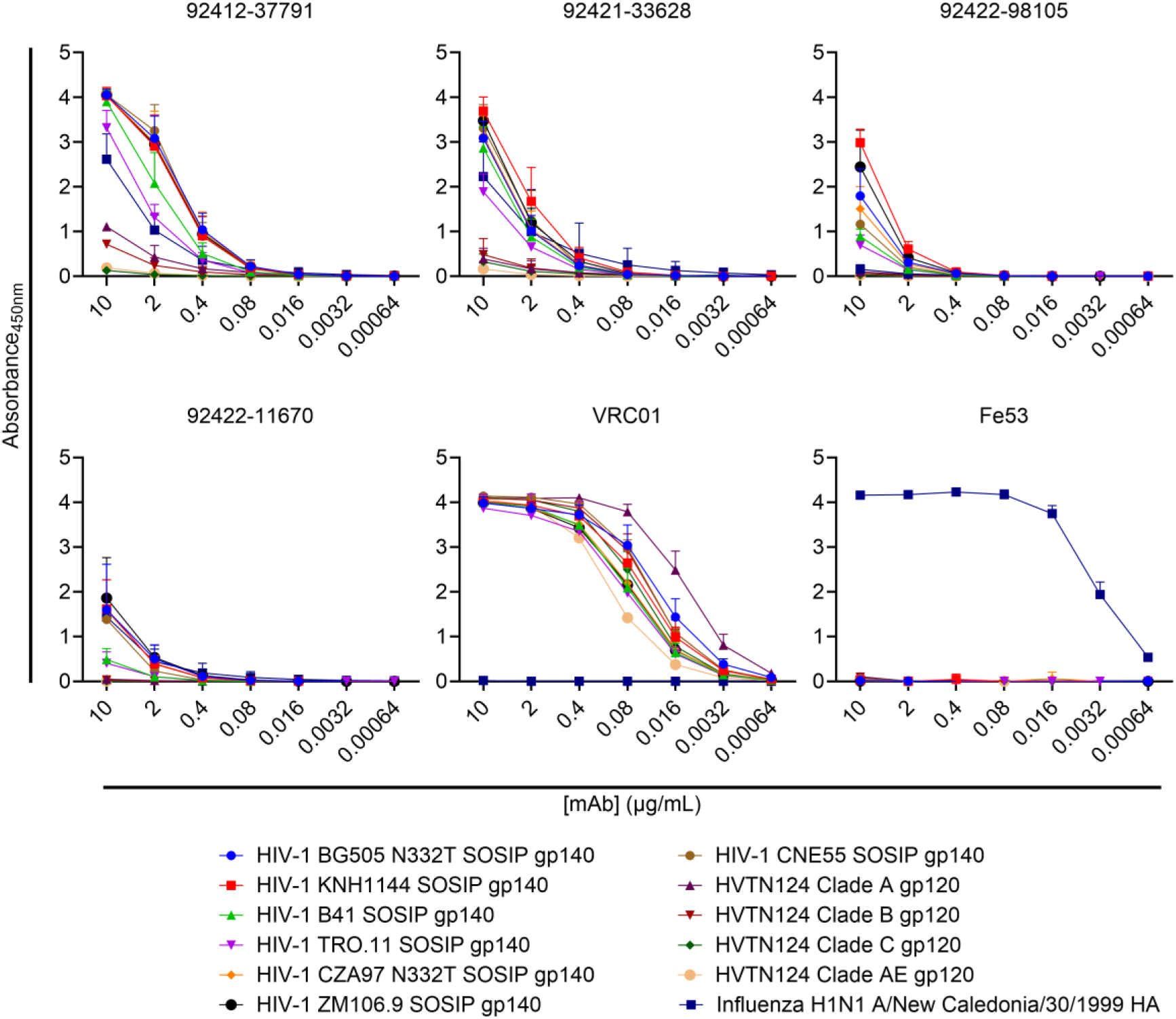
Characterization of HVTN124 mAbs by ELISA. ELISA validation of HVTN124 mAbs displayed as full curves. ELISA 5-fold curves from a set of three repeats in duplicate are displayed for each of the four HVTN124 mAbs, along with the VRC01 and Fe53 mAb controls, against all 12 antigens used in the LIBRA-seq screening library.

**Figure S2.**
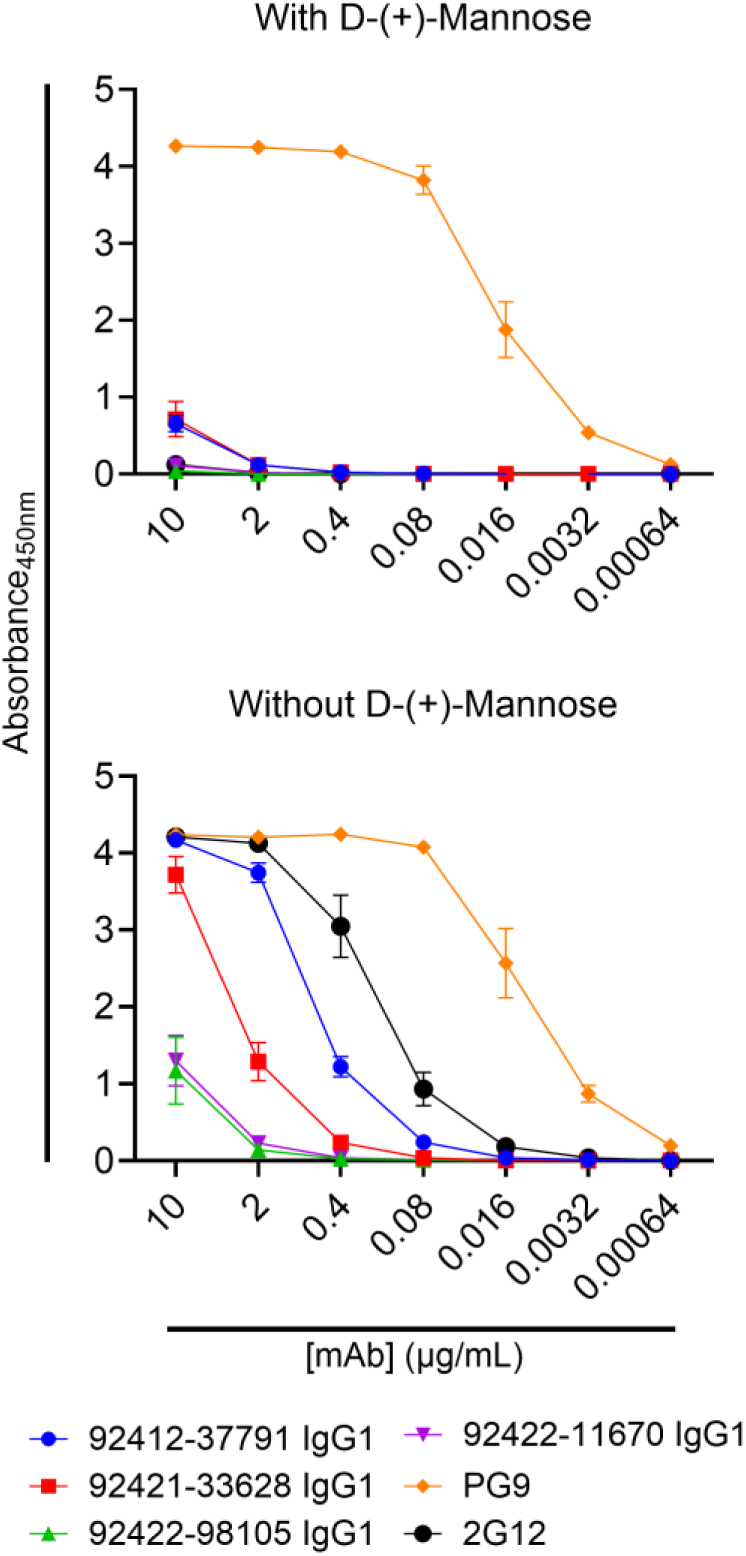
HVTN124 mAbs achieve broad reactivity via N-linked glycan recognition. Antibody competition with and without 1M D-(+)-Mannose displayed as full curves. Absorbance at 450nm is listed on the Y-axis, while antibody concentration in µg/mL is listed on the X-axis. The four HVTN124 mAbs, along with the V3-glycan-reactive 2G12 and V1/V2-reactive PG9 control mAbs, were incubated with and without 1M D-(+)- Mannose against HIV-1 CNE55 SOSIP gp140. ELISA 5-fold curves for both conditions from a set of three repeats in duplicate are displayed.

**Figure S3.**
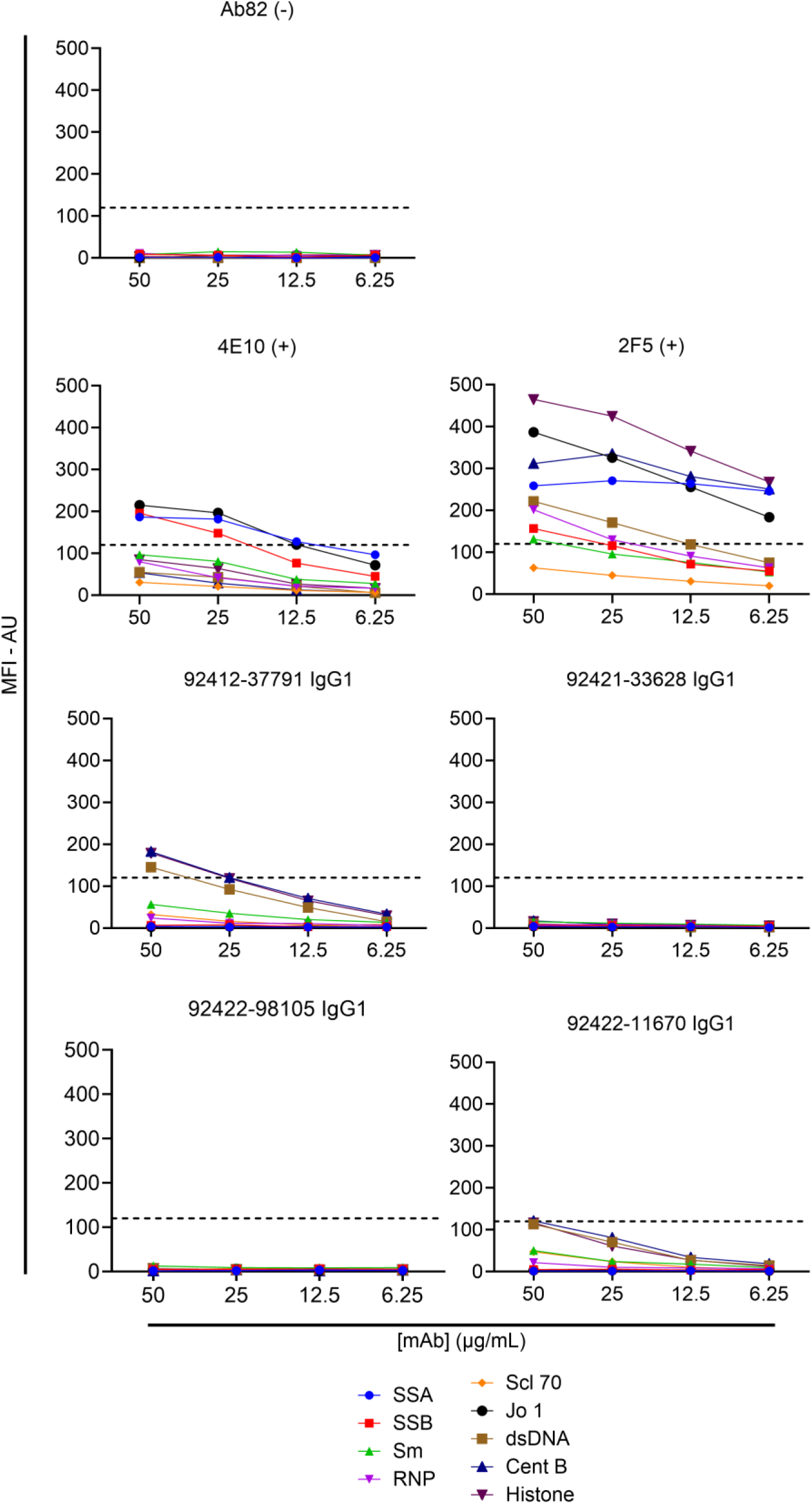
AtheNA autoreactivity analysis of HVTN124 mAbs. Autoantigen reactivity against the AtheNA panel as full curves. HVTN124 mAbs were tested for autoreactivity against the AtheNA panel. MFI – AU is listed on the Y-axis, while antibody concentration in µg/mL is listed on the X-axis. Positive control antibodies included 4E10 and 2F5, while Ab82 was used as a negative control antibody. Values exceeding 120 MFI at 25 µg/mL for the AtheNA assay are considered positive. Negative MFI – AU values were transformed to zero.

**Figure S4.**
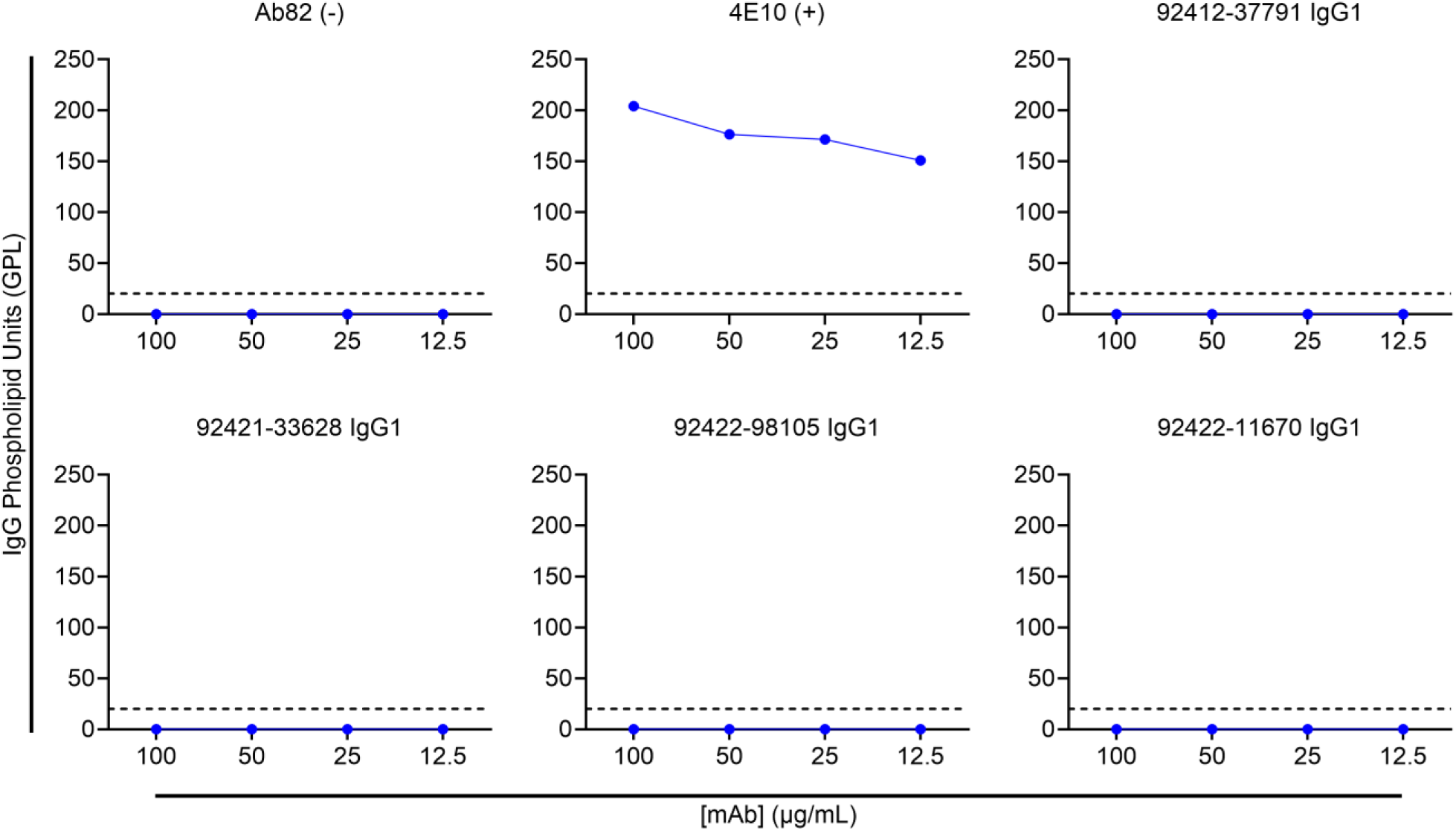
Cardiolipin autoreactivity analysis of HVTN124 mAbs. Autoreactivity toward cardiolipin as full curves. IgG phospholipid units (GPL) is listed on the Y-axis, while antibody concentration in µg/mL is listed on the X-axis. 4E10 was used as a positive control antibody, while Ab82 was used as a negative control antibody. Values at or greater than 20 GPL at 50 µg/mL are considered positive. Negative GPL values were transformed to zero.

**Figure S5.**
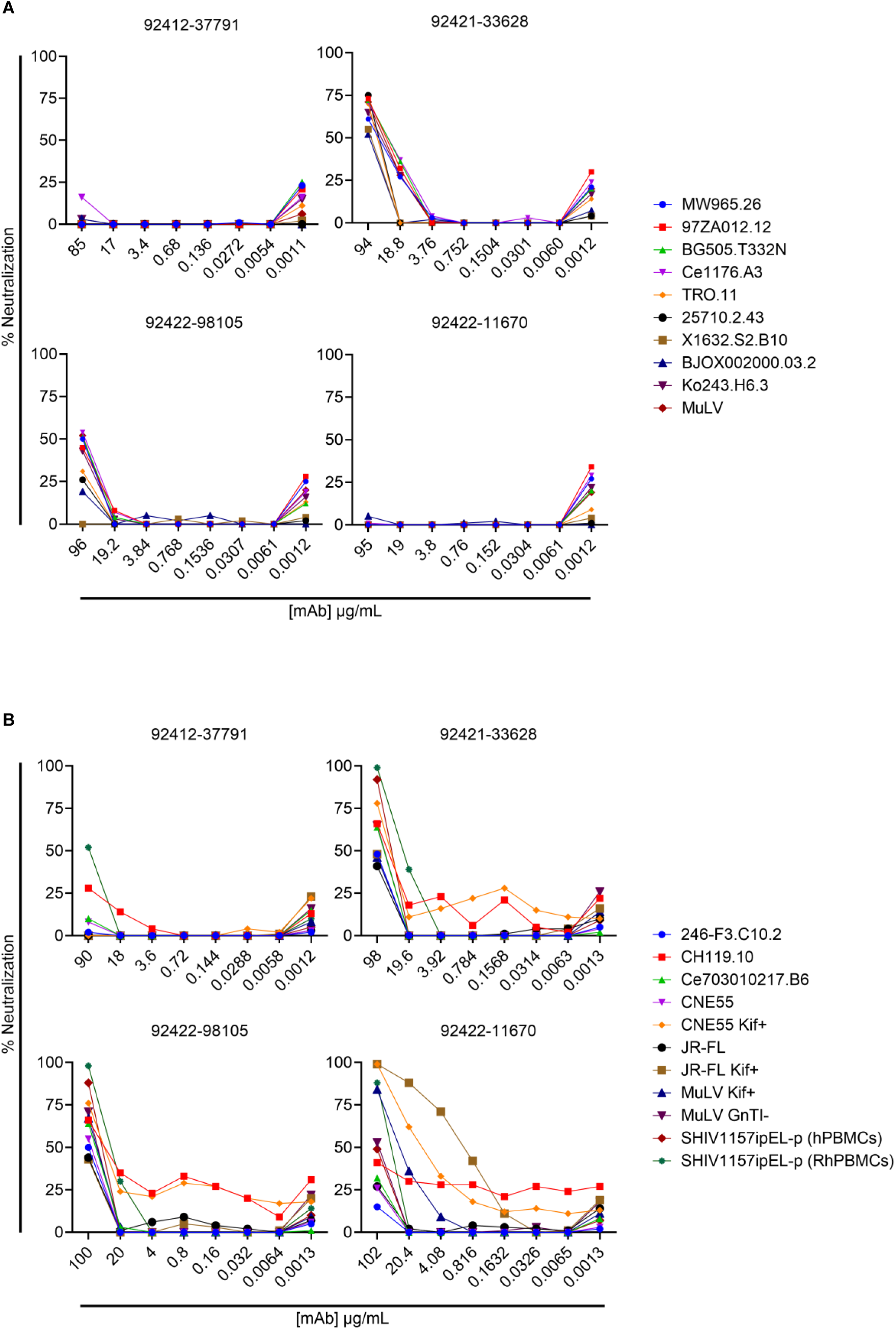
HIV-1 neutralization. (A) HVTN124 mAbs vs. HIV-1 pseudovirus set 1. Lot 1 of HVTN124 mAbs vs. HIV-1 pseudovirus. Percent neutralization is listed on the Y-axis while antibody concentration in µg/mL is listed on the X-axis. Percent neutralization of the 5-fold curves are displayed for each of the four HVTN124 mAbs being tested against: MMW965.26, 97ZA012.12, BG505.T332N, Ce1176.A3, TRO.11, 25710.2.43, X1632.S2.B10, BJOX002000.03.2, Ko243.H6.3, and MuLV. Negative percent neutralization values were transformed to zero. (B) HVTN124 mAbs vs. HIV-1 pseudovirus set 2. Percent neutralization is listed on the Y-axis while antibody concentration in µg/mL is listed on the X-axis. Percent neutralization of the 5-fold curves are displayed for each of the four HVTN124 mAbs being tested against: 246-F3.C10.2, CH119.10, Ce703010217.B6, CNE55, CNE55 Kif+, JR-FL, JR-FL Kif+, MuLV Kif+, MuLV GnTI-, SHIV1157ipEL-p (hPBMCs), and SHIV1157ipEL-p (RhPBMCs). Negative percent neutralization values were transformed to zero.

**Figure S6.**
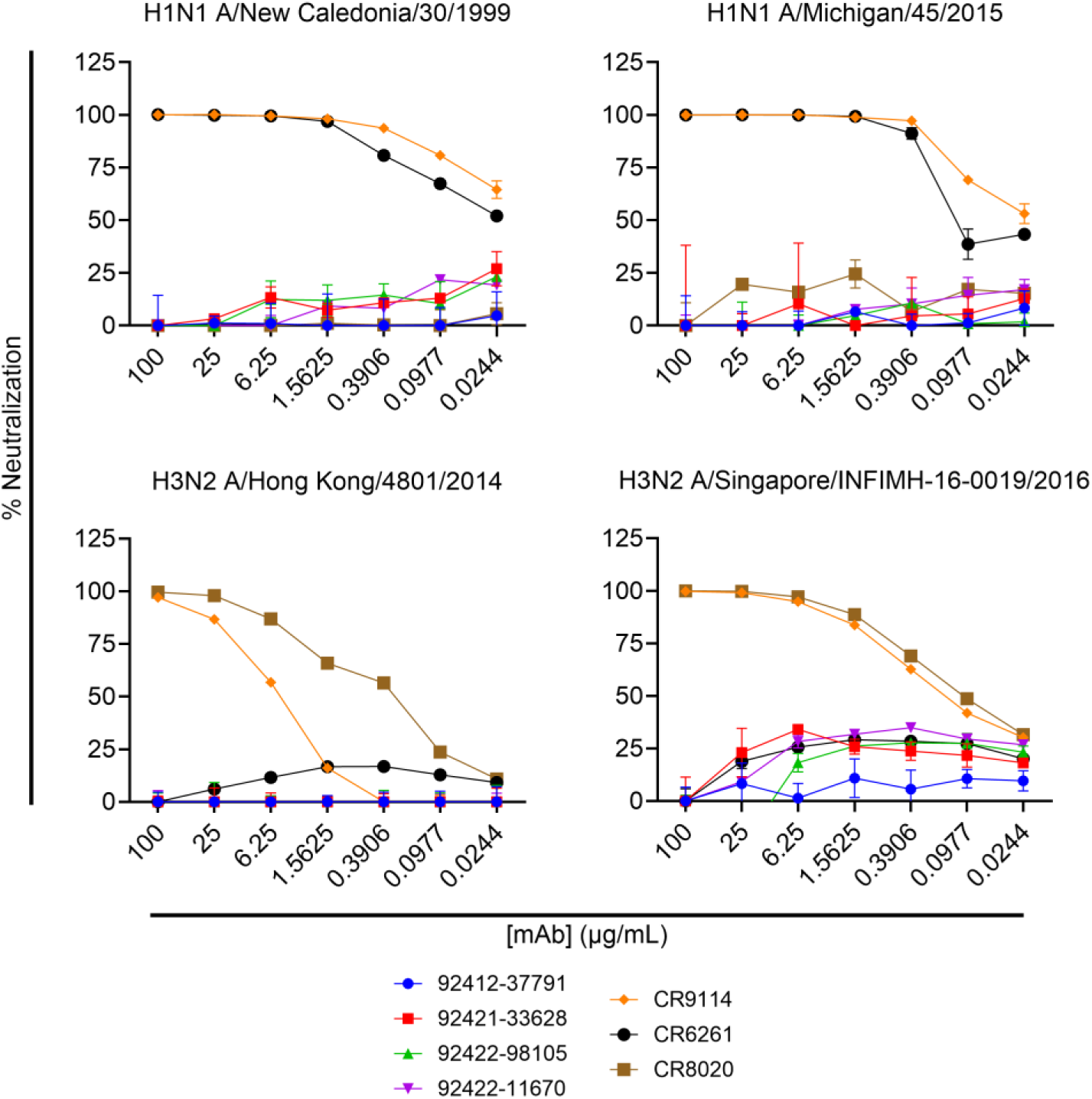
Influenza neutralization. Neutralization of HVTN124 mAbs vs. H1N1 and H3N2 Influenza viruses as 4-fold curves. HVTN124 mAbs were tested for neutralizing activity against an influenza viral panel that included: H1N1 A/New Caledonia/30/1999, H1N1 A/Michigan/45/2015, H3N2 A/Singapore/INFIMH-16-0019/2016, and H3N2 A/Hong Kong/4801/2014. Of the control mAbs used, CR9114 is known to broadly neutralize influenza A and B viruses, while CR6261 targets group 1 influenza A viruses such as H1N1 isolates, and CR8020 targets group 2 influenza A viruses such as H3N2 isolates. The percent neutralization is listed on the Y-axis, while the antibody concentrations in μg/mL is listed on the X-axis. Negative percent neutralization values were transformed to zero.

**Figure S7.**
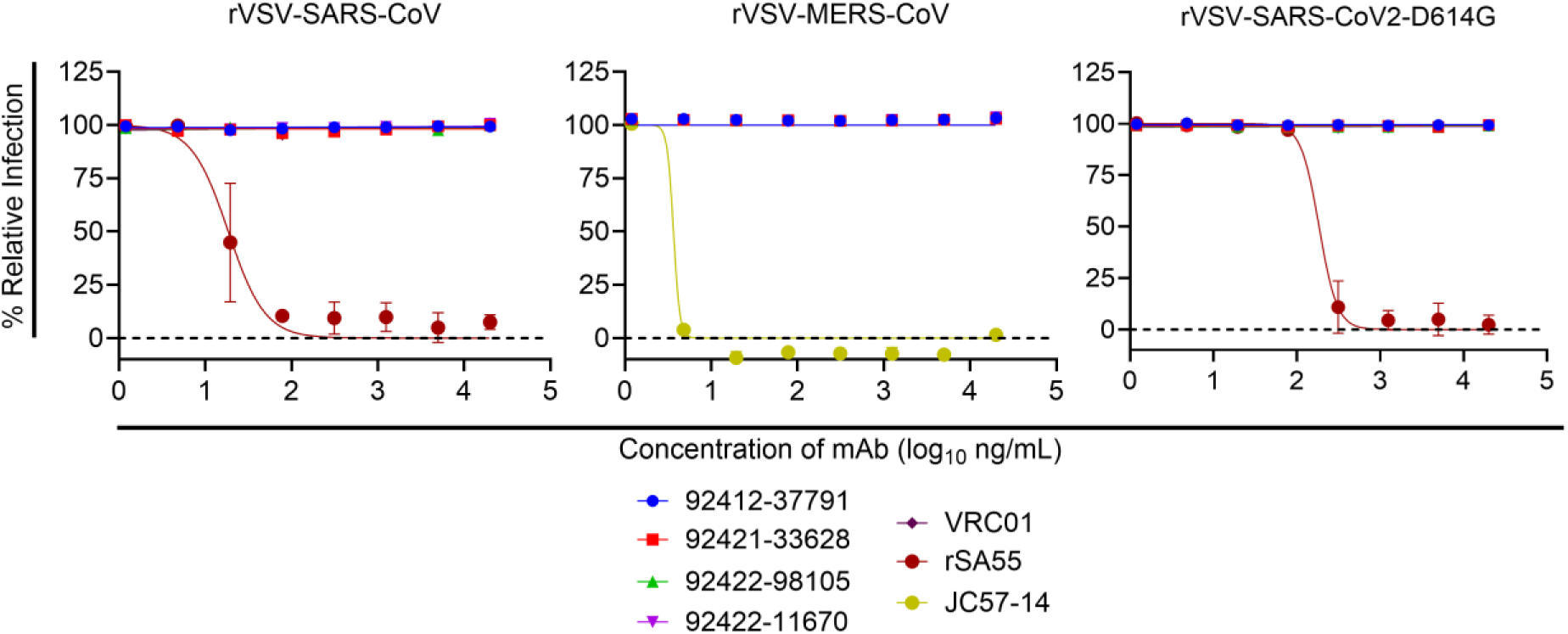
Coronavirus neutralization. Coronavirus neutralization as 4-fold curves. HVTN124 mAbs were screened for neutralizing activity against rVSV-SARS-CoV-2 D614G, rVSV-MERS-CoV, and rVSV-SARS-CoV. Positive controls included rSA55 for rVSV-SARS-CoV-2 D614G and rVSV-SARS-CoV, while JC57-14 was used for rVSV-MERS-CoV. VRC01 was also included as a negative control mAb. The percent infectivity is listed on the Y-axis, while the antibody concentrations in μg/mL mAb is listed on the X-axis. Data are mean ± standard deviations (SD) of technical triplicates from a representative experiment repeated twice.

